# A constricted mitochondrial morphology optimizes respiration

**DOI:** 10.1101/2024.03.21.582105

**Authors:** Manish K. Singh, Laetitia Cavellini, Christina Kunz, Mickaël Lelek, Perrine Bomme, Naïma Belgareh-Touzé, Adeline Mallet, Lea Dietrich, Christophe Zimmer, Mickael M. Cohen

## Abstract

Mitochondria assemble in a dynamic tubular network with a morphology governed by mitochondrial fusion and fission, which regulate all mitochondrial functions including oxidative phosphorylation ^1–4^. Yet, the link between mitochondrial morphology and respiration remains unclear ^5–9^. Here, we discover a previously unknown mitochondrial morphology dedicated to respiratory growth of *Saccharomyces cerevisiae*, which we refer to as “Ringo”. The Ringo morphology is characterized by stable constrictions of mitochondrial tubules. Ringo constrictions are mediated by the yeast dynamin Dnm1 and, unlike mitochondrial fission ^10–12^, occur in the absence of contacts with the endoplasmic reticulum. Our data show that the Ringo morphology regulates mitochondrial DNA homeostasis during respiration to ensure stoichiometric assembly of OXPHOS complexes, demonstrating that the shape of mitochondria actively contributes to optimal respiration.

**One-Sentence Summary:** We report a new mitochondrial morphology that actively contributes to optimal respiration in yeast.

## Main Text

Mitochondria form tubular structures inside the cell that associate in an intricate membrane network. The morphology of these tubules is conditioned by an equilibrium between frequent fusion and fission events mediated by large GTPases of the Dynamin-Related Proteins (DRP) superfamily ^1–3^. Among the DRPs, Dnm1 and DRP1 mediate fission of the outer mitochondrial membrane in yeast and metazoans ^2,4,12,13^. The DRPs are recruited to mitochondrial membranes by specific mitochondrial adaptors that accumulate at sites of contact between mitochondria and the Endoplasmic Reticulum (ER). Once recruited, Dnm1 and DRP1 auto-oligomerize in a GTP-dependent manner to form macromolecular spirals which wrap around mitochondrial tubules that have been pre-constricted by the ER ^10,13–15^. GTP hydrolysis reduces the diameter of the spiral, resulting in hyper-constriction of the mitochondrial tubule and eventual fission. Together, mitochondrial fusion and fission guide mitochondrial dynamics and regulate key mitochondrial functions including oxidative phosphorylation ^1,5,6^. Yet, it is not known to what extent the morphology of the mitochondrial network affects respiration ^7^. The model organism *Saccharomyces cerevisiae* is an ideal system to study the crosstalk between mitochondrial dynamics and metabolism. Depending on the carbon source that is provided in the culture medium, *Saccharomyces cerevisiae* can grow either through respiration or fermentation. Glycerol or ethanol induce respiratory growth, whereas D-glucose, also called dextrose, inhibit respiration even in the presence of oxygen, and is a preferred fermentative carbon sources ^16,17^.

### Discovery of the Ringo mitochondrial morphology

We analysed mitochondrial morphology during fermentation (dextrose containing media, YPD) or respiration (glycerol containing media, YPG) first using confocal microscopy in cells in which the outer mitochondrial membrane (Tom70-GFP) and matrix (mito-mCherry) were fluorescently labelled. In confocal images, whose resolution is ∼250 nm, mitochondrial outer membranes and matrices were co-localized and no differences in morphology were observed during fermentation compared with respiration (Fig. 1a). Structured Illumination Microscopy (SIM) is a super-resolution imaging method that improves the resolution to ∼125 nm. Thanks to the improved resolution of SIM, mitochondria in fermentation conditions clearly appeared as tubes, with a thin tubular matrix surrounded by the outer membrane (Fig. 1b and S1a). Strikingly, SIM revealed that tubules underwent extensive rearrangements upon respiration. This reorganization is reflected in numerous constrictions of the outer membrane, which resemble beads on a string in 2D. The matrix, on the other hand, appeared disconnected (Fig. 1b and S1a). We name this morphology of respiratory mitochondria, which to our knowledge has not been reported before, “Ringo”.

**Fig. 1.**
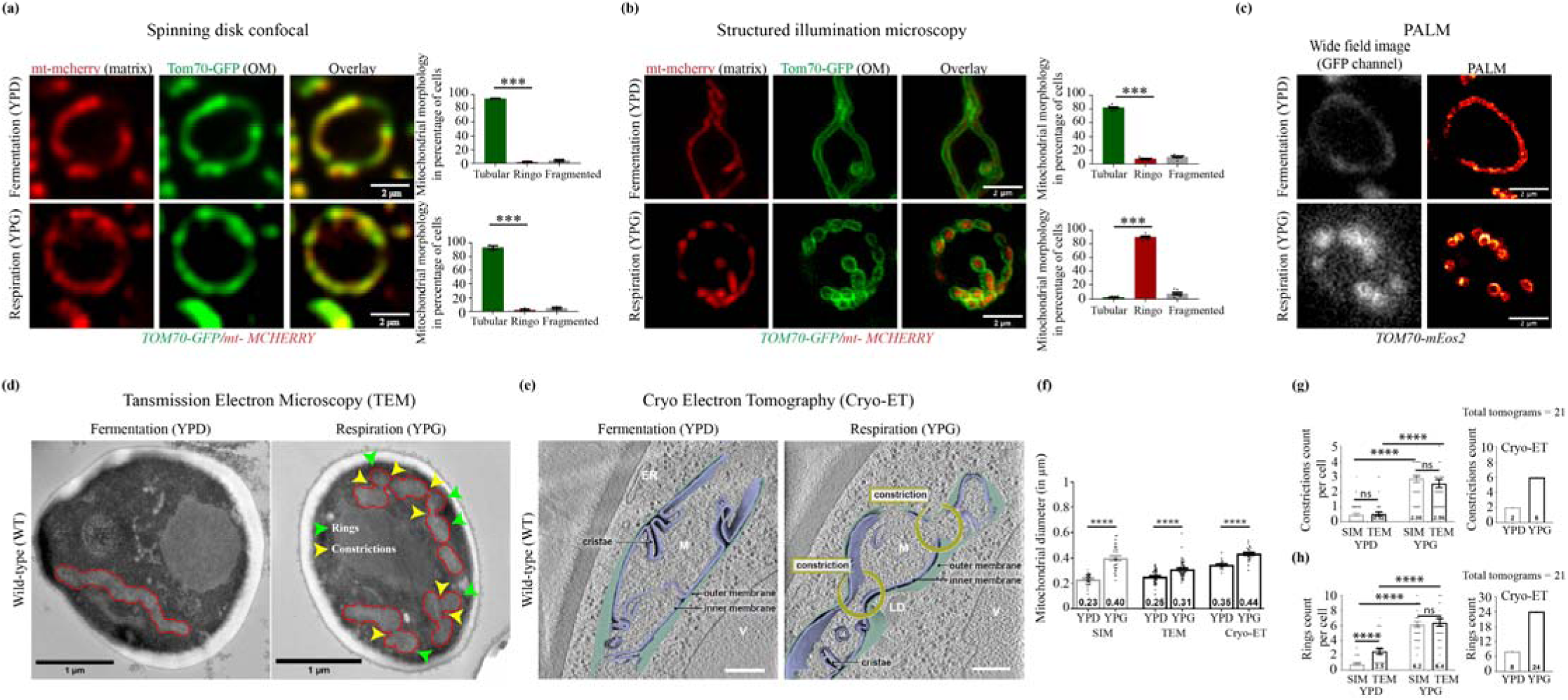
Identification and characterization of the Ringo mitochondrial morphology. **(a)** Spinning-disc confocal acquisitions of cells labeled for mitochondrial matrix (mt-mCherry) and Outer Membranes (Tom70-GFP) in fermentation (top) or respiration (bottom). Scale bar, 2 µm. Right graphs: percentage of cells with Tubular, Ringo or Fragmented mitochondria. Mean ± SEM from >192 cells in 3 independent experiments. **(b)** Same as (a) but in SIM and quantifications with >329 cells. **(c)** Wide-field or PALM images of cells labeled with TOM70-mEOS2 in fermentation or respiration. **(d)** TEM micrographs of cells in fermentation or respiration (Scale bar,1 µm). Mitochondria are delimited by red demarcations. Rings and constrictions are indicated by green or yellow arrowheads, respectively. **(e)** Slices through tomographic volumes and 3D renderings of mitochondria (M) during fermentation or respiration. Vacuoles (V), Lipid Droplets (LD) and mitochondrial sub-compartments are indicated. Scale bars, 500 nm. **(f)** Average mitochondrial diameter, **(g)** constriction counts per cell or per Tomogram and **(h)** ring counts per cell or per Tomogram, in YPD or YPG as quantified in SIM, TEM, or Cryo-ET. Mean ± SEM from 50 (SIM), 48 (TEM) and 21 (Cryo-ET) images. *P<0.05, **P<0.01, ***P<0.001, ****P<0.0001, ns, not significant (Mann-Whitney t-test).

To verify this finding with a different imaging method, we used photo activated localization microscopy (PALM), which achieves spatial resolutions of 20-50 nm, in Tom70-mEos2 cells. Mitochondrial networks imaged by PALM in cells grown in dextrose (YPD) were tubular, while those imaged in cells grown in glycerol (YPG) were formed of connected rings similar to those captured by SIM, providing independent confirmation of the Ringo morphology (Fig. 1c and S1b). The Ringo morphology was observed regardless of the protein used to label the outer mitochondrial membrane (either Om45-GFP or Tom70-GFP), indicating that this phenotype is a property of the outer membrane shape rather than of specific membrane proteins (Fig. 1b and Extended Data Fig. 1c).

### The Ringo mitochondrial morphology is linked to respiration

Our clear-cut observation that the Ringo morphology occurs during cell growth in glycerol media while the mitochondrial network is essentially tubular in dextrose media suggests a possible functional link between the formation of Ringo mitochondrial networks and oxidative phosphorylation. To further test this association, we analysed mitochondrial morphology in ethanol, another medium that promotes respiration. Again, the Ringo morphology was dominant (Extended Data Fig. 2a). When yeast is grown in dextrose media, this source of carbon is progressively depleted, and cells switch to self-produced ethanol as a respiratory carbon source to continue growing. If the Ringo morphology is associated with respiration, we therefore expect a switch from a tubular morphology to the Ringo morphology as function of time. We indeed observe this switch after 4-6 hours of growth in 2% dextrose media (Extended Data Fig. 2b). If this switch is due to dextrose depletion, it should occur earlier if less dextrose is available initially. This prediction is borne out in experiments where we lowered the percentage of dextrose (Fig. S2c). Conversely, we confirmed that the Ringo morphology reverts to the classical tubular morphology when cells get transferred from glycerol to dextrose media (Extended Data Fig. 2d). Unlike in glycerol or ethanol, cells grown in galactose or raffinose media can use both respiratory and fermentative growth ^18–20^. If Ringo and tubular morphologies are associated with respiration and fermentation, respectively, we expect a mixture of these morphologies. Indeed, we observed that when grown in media with either of these two distinct carbon sources, 27% of cells displayed tubular mitochondria whereas 63% had Ringo mitochondrial networks (Extended Data Fig. 2e). These experiments further support a close link between the Ringo morphology and respiration.

### The Ringo morphology is formed by constrictions of the mitochondrial network

Axial projections of cells expressing Tom70-GFP (outer membrane) and mito-mCherry (matrix) as mitochondrial markers of tubular (Extended Data Fig. 3a), fragmented (Extended Data Fig. 3b) or Ringo networks (Extended Data Fig. 3c), suggested that mitochondria with Ringo morphology are not separated from each other (Extended Data Fig. 3c). Instead, the matrix apparently remains continuous between each outer membrane compartment of the Ringo network (Extended Data Fig. 3c), suggesting an unconventional mitochondrial architecture maintained by repetitive constrictions of mitochondrial tubules. However, the resolution of SIM or PALM are insufficient to rule out discontinuous membranes in close proximity resulting from a fission event. We therefore complemented fluorescence imaging with Transmission Electron microscopy (TEM) and Cryo-Electron Tomography (Cryo-ET) analysis. These methods confirmed that mitochondria of cells grown in dextrose-containing media (YPD) were overall tubular, whereas mitochondria of cells grown in glycerol-containing media (YPG) were remodelled by numerous constrictions (Fig. 1d-e and Extended Data Fig. 4-5). The diameter (Fig. 1f) of Ringo mitochondria (0.31 to 0.44 μm) was wider than that of tubular mitochondria (0.23 to 0.35 μm). In glycerol as compared to dextrose media, the number mitochondrial constrictions (Fig. 1g) increased by 3 (Cryo-ET) to 6 (SIM and TEM) fold and the number of rings (Fig. 1h) by 2 (TEM), 3 (cryo-ET) or 6 (SIM) fold. These results thus confirm major changes in mitochondrial morphology upon respiratory (YPG) as compared to fermentative growth (YPD) and reveal that the Ringo morphology is formed by numerous constrictions of mitochondrial tubules.

### Dnm1 and Mdv1 are required for formation of the Ringo morphology

Mitochondrial constriction is generally considered to be a prerequisite for fission, which in yeast involves the Dnm1 DRP ^2,4,12,13^. If Dnm1 also promotes constrictions in the Ringo morphology, its ablation should abolish formation of this phenotype. SIM analysis of *dnm1Δ* cells labelled with Tom70-GFP and mito-mCherry confirmed that the absence of Dnm1 results in hyperfused mitochondria ^12^ upon fermentative growth (Fig. 2a and S6a, YPD). Respiratory conditions that produce the Ringo morphology in *WT* cells, led to another previously unknown mitochondrial phenotype in *dnm1Δ* cells (Fig. 2a and S6a, YPG). The *dnm1Δ* mitochondrial network is characterized by significantly wider mitochondrial diameters and equally frequent but seemingly more random constrictions than in the Ringo morphology (Extended Data Fig. 6b). We named this new mitochondrial morphology Muskaan (after the newborn daughter of M. K. Singh).

**Fig. 2.**
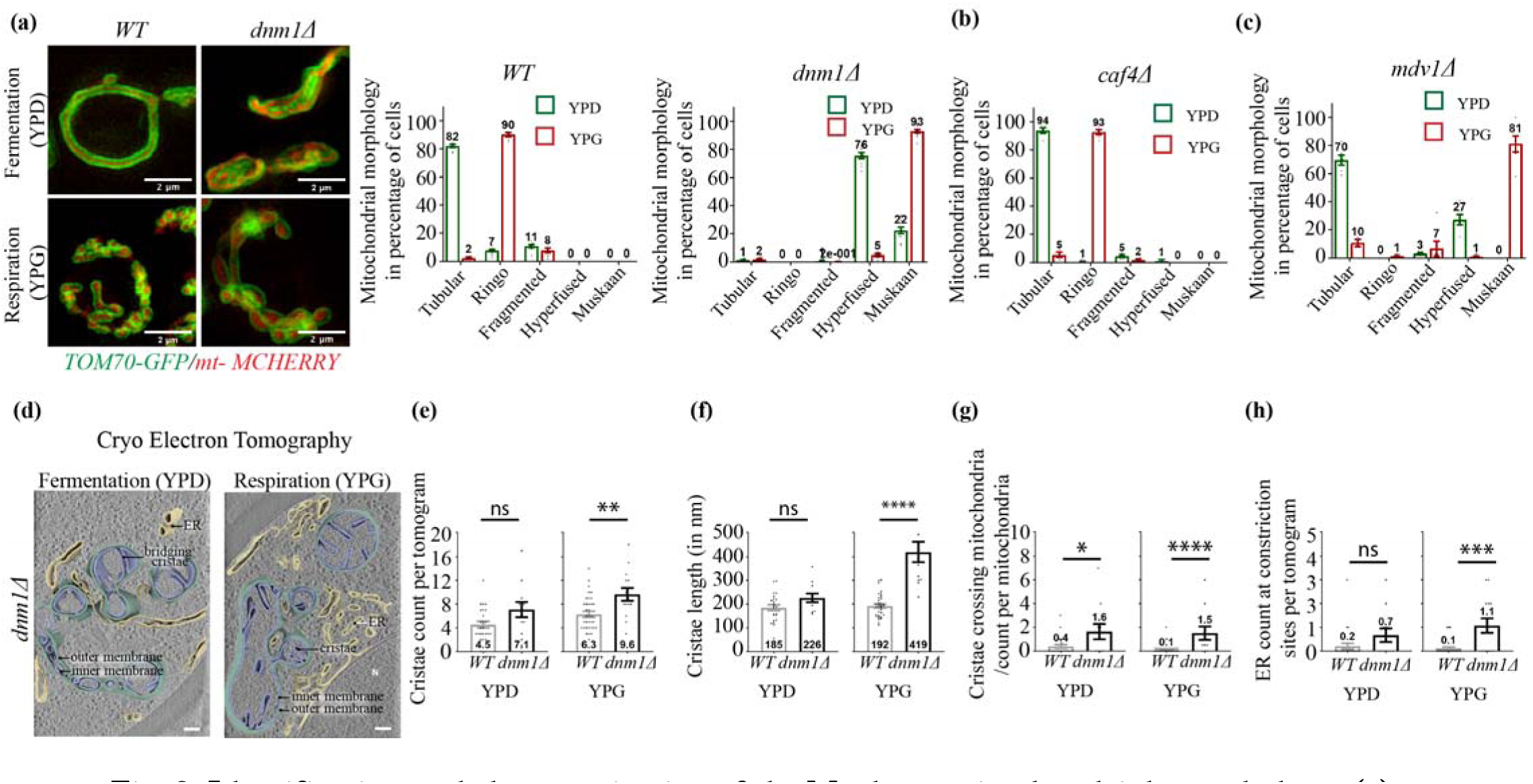
Identification and characterization of the Muskaan mitochondrial morphology. **(a)** SIM acquisitions of *WT* or *dnm1*Δ cells labeled for mitochondrial matrix (mt-mCherry) and Outer Membranes (Tom70-GFP) in fermentation (top) or respiration (bottom). Scale bar, 2 µm. Right graphs: percentage of cells with Tubular, Ringo, Fragmented, Hyperfused or Muskaan mitochondria during fermentation (YPD, green) or respiration (YPG, red). Mean ± SEM from >178 cells in 3 independent experiments. **(b and c)** Same as graphs from (a) in *caf4*Δ (b) and *mdv1*Δ (c) cells. **(d)** Slices through tomographic volumes and 3D renderings of Muskaan mitochondria from *dnm1*Δ cells during fermentation or respiration. Endoplasmic Reticulum (ER), Nucleus (N) and mitochondrial sub-compartments are indicated. Scale bars, 500 nm. **(e)** Cristae count per tomogram, **(f)** Cristae length, **(g)** Cristae crossing mitochondria count per mitochondria and **(h)** ER count at constrictions sites per tomogram in *WT* or *dnm1*Δ during fermentation (YPD) and respiration (YPG), as quantified in Cryo-ET. Mean ± SEM from 21 (*WT*) or 11 (*dnm1*Δ) tomogram. *P<0.05, **P<0.01, ***P<0.001, ****P<0.0001, ns, not significant (Mann-Whitney t-test).

Mitochondrial recruitment of Dnm1 occurs through the mitochondrial adaptors Mdv1 or Caf4 which can themselves interact with the mitochondrial anchor Fis1 ^21–26^. Absence of Caf4 neither affected tubular morphology in fermentation nor the Ringo morphology in respiration (Fig. 2b and S6c). In *mdv1Δ* cells, however, the Ringo morphology was no longer observed in respiratory growth and was instead replaced by the Muskaan morphology (Fig. 2c and S6d). These results indicate that proper formation of the Ringo phenotype requires Dnm1 and Mdv1 but not Caf4.

The Muskaan morphology captured in the absence of Dnm1 or Mdv1 is clearly distinct from the Ringo morphology seen in *WT* cells. Cryo-ET analysis further revealed that mitochondria from *dnm1Δ* cells grown in respiratory conditions display a significant reorganization of cristae as compared to mitochondria from *WT* cells (Fig. 2d and 2e). Cristea of Muskaan mitochondria were not only much longer than those of Ringo networks (Fig. 2f) but also often extended across the whole width of the organelle (Fig. 2g), leading to numerous separations of the mitochondrial matrix (Fig. 2d, bridging cristae). Another intriguing difference between Muskaan and Ringo networks relates to the presence of ER at constriction sites (Fig. 2h). Constriction sites of hyperfused and Muskaan mitochondria were systematically decorated with ER membranes, which are known to be required to initiate mitochondrial fission ^10^. This is consistent with abortive mitochondrial division in the absence of Dnm1. In contrast, the ER was only rarely detected in the vicinity of the numerous constrictions of Ringo networks (Fig. 2h). These results suggest that Ringo constrictions require Dnm1 but not contact sites with the ER.

### The Ringo morphology is formed in the absence of contacts between mitochondria and the Endoplasmic Reticulum

In yeast, contact sites between the ER and mitochondria are mediated by the ERMES (Endoplasmic Reticulum and Mitochondria Encounter Structures) complex that includes four core components – Mmm1, Mdm34, Mdm12 and Mdm10 - with Mmm1 being integral to ER membranes ^11^. As a consequence, mitochondrial localization of Mmm1 signals contact sites with the ER. To further assess the roles of Dnm1 and the ER, we analysed the localization of Dnm1-GFP and Mmm1-GFP on mitochondria labelled with Tom70-mCherry under fermentative and respiratory conditions (Fig. 3a; left images). Surprisingly, we observed a two-fold increase in the number of Dnm1-GFP puncta upon respiratory compared to fermentative growth (Fig. 3a; right graphs). This difference was confirmed by western blot analysis, in which the expression of endogenous Dnm1 increased significantly upon respiratory growth (Fig. 3b; left blot). The increase in Dnm1 is transcriptionally regulated, since placing *DNM1* expression under the control of the constitutive TEF promoter abolished Dnm1 variations between fermentation and respiration (Fig. 3c). Most importantly, in contrast to Dnm1, no changes in the number of Mmm1-GFP puncta or the expression levels of Mmm1-GFP were observed (Fig. 3a, right graph and 3b, right blot). Consistent with this, Dnm1 recruitment on mitochondria increased by more than two-fold in respiration as compared to fermentation, whereas Mmm1 was unchanged (Fig. 3d).

**Fig. 3.**
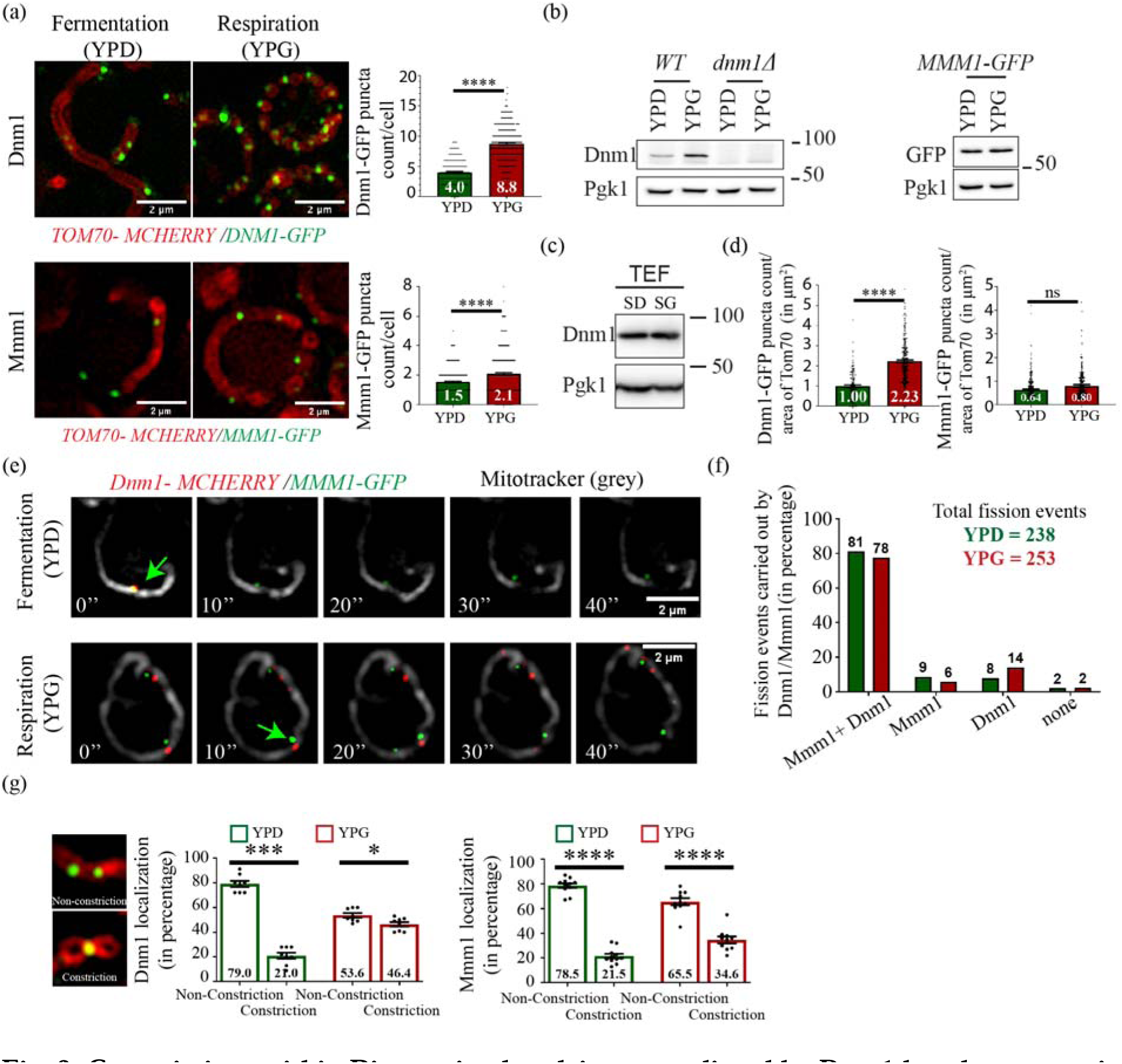
Constrictions within Ringo mitochondria are mediated by Dnm1 but do not require contacts with the ER. **(a)** SIM of *WT* cells labeled for Tom70-mcherry and Dnm1-GFP (top) or Mmm1-GFP (bottom) in fermentation (YPD) or respiration (YPG). Scale bar, 2 µm. Right graphs: Dnm1 (top) and Mmm1 (bottom) puncta count per cell (>198) in YPD (green) or YPG (red). **(b)** Total protein extracts prepared from *WT* and *dnm1*Δ (left) or *MMM1-GFP* (right) cells in YPD or YPG and analyzed by immunoblotting as indicated. MW (kDa) indicated on the right. **(c)** Same as (b) with *TEF-DNM1* cells. **(d)** Quantification of Dnm1-GFP (left) or Mmm1-GFP (right) puncta count/area of Tom70 from cells (>196) in (a). **(e)** SIM Time-lapse from Dnm1-mcherry, Mmm1-GFP, mitotracker (grey) triple-labeled cells during fermentation and respiration. Green arrows indicate fission events. **(f)** Percentage of total fission events positive or negative for both Dnm1 and/or Mmm1. **(g)** Percentage of Dnm1 (left) or Mmm1 (right) localization at non-constrictions or constriction sites in cells from (a). For all graphs, Mean ± SEM and 3 independent experiments. *P<0.05, **P<0.01, ***P<0.001, ****P<0.0001, ns, not significant (Mann-Whitney t-test).

We reasoned that the mitochondrial recruitment of Dnm1 without ER contacts generates stable constrictions as those seen in the Ringo network whereas Dnm1 recruitment at mitochondria-ER (mito-ER) contact sites leads to mitochondrial fission. To test this hypothesis, we followed mitochondria fission over time using SIM (Extended Data Fig. 7a) and assessed co-recruitment of Mmm1 and Dnm1 at mitochondrial fission sites (Fig. 3e-f). This experiment revealed that the rate of mitochondrial fission is identical in fermentation (where tubular mitochondria dominate) and in respiration (where Ringo networks dominate), with on average 0.28 fission events per cell over 3 minutes of acquisition (Extended Data Fig. 7a). Quantification of Mmm1-GFP and Dnm1-mCherry on more than 200 fission sites also confirmed that both proteins co-localize in the vast majority (about 80%) of fission events (Fig. 3f). These results confirm that Dnm1 recruitment at mito-ER contact sites correlate with effective mitochondrial fission. In contrast, Dnm1 recruitment at mitochondria but not at mito-ER contacts would generate regular constrictions forming the Ringo networks. We evaluated this further by quantifying the amount of Dnm1 and Mmm1 puncta at mitochondrial constriction and non-constriction sites (Fig. 3g left images). Importantly, we observed that Dnm1 localization at non-constriction sites is significantly lower in respiration than in fermentation, and its localization at the junction of rings is more than double in respiration than in fermentation (Fig. 3g, left graphs). In contrast, no such redistribution between fermentation and respiration was observed for Mmm1 (Fig. 3g, right graphs). These results support a mechanism in which Dnm1 triggers repetitive constrictions within the Ringo mitochondrial network that do not result in fission in the absence of contacts with the ER.

### The Ringo morphology is required for optimal respiration

Cells that respire need an increased mitochondrial protein import than cells that undergo fermentation ^27^. We therefore predict that in conditions that allow for the presence of both morphologies, Ringo networks are associated with enhanced import than tubular networks. To test this prediction, we used a preCox4-mCherry construct as a marker for mitochondrial import efficiency ^28^ and performed dual color imaging with Tom70-GFP upon growth in galactose and raffinose media. We observed that mitochondrial mCherry intensity is 30% to 35% higher in cells with Ringo than cells with tubular networks (Extended Data Fig. 7b), thereby providing yet another confirmation that the Ringo phenotype is associated with respiration.

To explore the causal relationships between the Ringo morphology and respiration, we next tested if blocking oxidative phosphorylation would prevent the Ringo phenotype. Treatment of *WT* cells with 1 μM Antimicyin A, an inhibitor of the respiratory chain acting on complex III ^29^, did not affect tubular morphology in dextrose media (Extended Data Fig. 8a, top graphs) but induced extensive fragmentation of the Ringo network in glycerol already 15 minutes after treatment (Extended Data Fig. 8a, bottom graphs). Notably, fragmented mitochondria gradually fused into tubular networks after 60 min of treatment but the Ringo morphology was never recovered in this timeframe (Extended Data Fig. 8a, bottom graphs). These results suggest that oxidative phosphorylation is essential for maintaining the Ringo morphology.

Conversely, we asked whether blocking the formation of Ringo networks affects respiration. For this purpose, we relied on our observation that absence of Dnm1 induces disorganization of the Ringo network, giving rise to the Muskaan morphology (Fig. 2). We observed that Muskaan mitochondria from *dnm1Δ* cells display a 23% decrease in preCox4-mCherry mitochondrial import as compared to Ringo networks from *WT* cells (Extended Data Fig. 8b; YPG; *WT* vs *dnm1Δ*). This decrease in protein import was accompanied by impaired growth of *dnm1Δ* cells on glycerol media (Fig. 4a; YPG). Thus, *DNM1* ablation results in significant inhibition of respiration. Notably, reintroduction of Dnm1 in *dnm1Δ* cells (Extended Data Fig. 8c) not only restored respiratory growth (Fig. 4a; YPG) but also reversed mitochondrial morphology from Muskaan to Ringo networks (Fig. 4b, S8d and S8e). The remaining aggregated, Muskaan and hyperfused mitochondria are likely caused by random loss of the *DNM1* plasmid from *dnm1Δ* cells (Fig. 4b, S8d and S8e). These results indicate that inhibition of both respiratory capacity and Ringo morphology formation are exclusively caused by the absence of Dnm1.

**Fig. 4.**
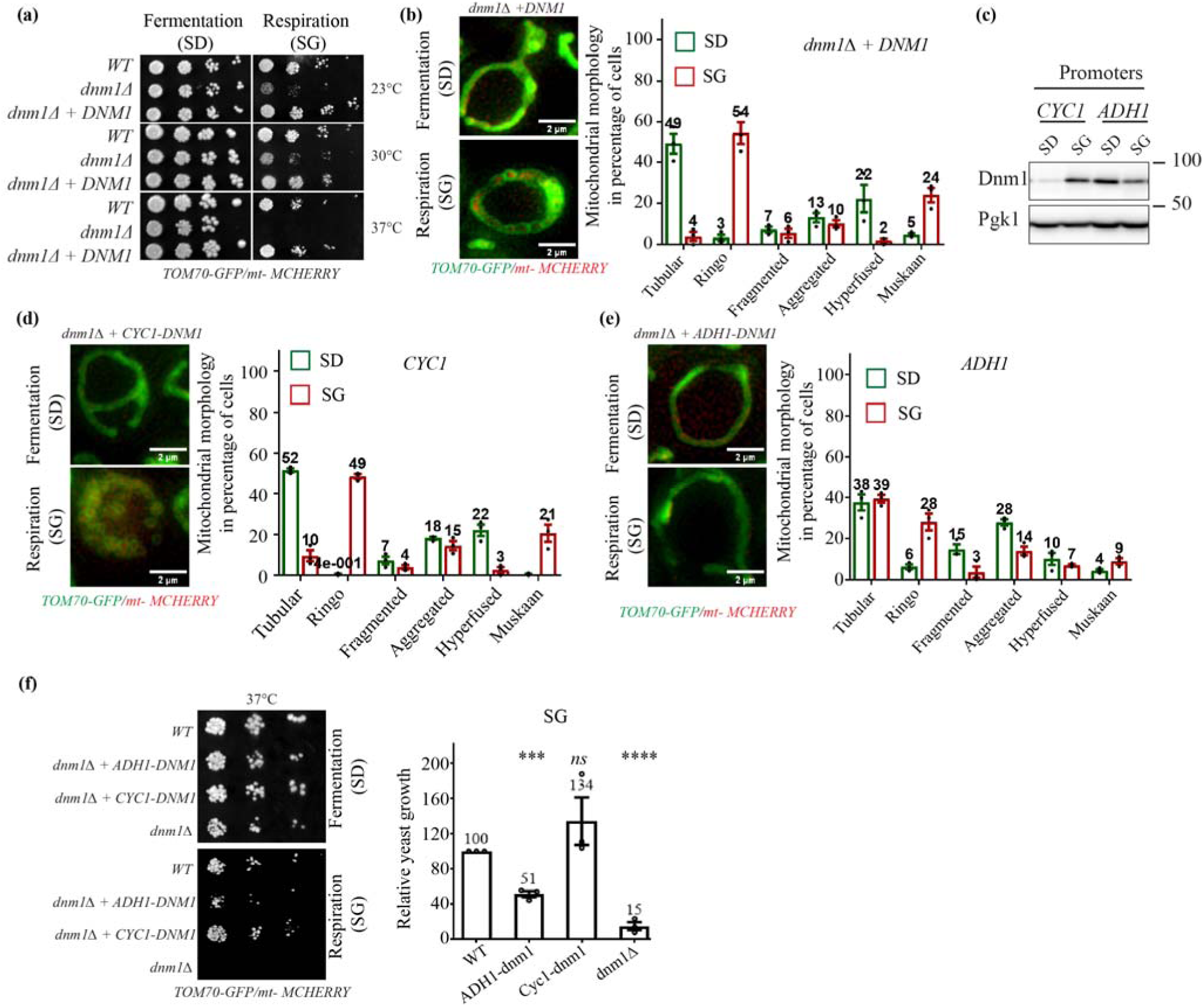
The Ringo morphology is required for optimal respiration. **(a)** Dextrose and glycerol serial dilutions of *WT*, *dnm1*Δ and *dnm1*Δ*+DNM1* strains at 23, 30 and 37^0^C. **(b)** SIM acquisitions of *dnm1*Δ*+DNM1* cells labeled with mt-mCherry and Tom70-GFP in fermentation (top) or respiration (bottom). Scale bar, 2 µm. Right graph: percentage of cells with Tubular, Ringo, Fragmented, Aggregated, Hyperfused or Muskaan mitochondria during fermentation (SD, green) or respiration (SG, red). Mean ± SEM from >198 cells in 3 independent experiments. **(c)** Total protein extracts prepared from *dnm1*Δ*+CYC1-DNM1* or *ADH1-DNM1* cells in dextrose (SD) or glycerol (SG) media and analyzed by immunoblotting as indicated. MW (kDa) indicated on the right. **(d)** Same as (b) with *dnm1*Δ*+CYC1-DNM1* cells (>207 cells). **(e)** Same as (b) with *dnm1*Δ*+ADH1-DNM1* cells (264 cells). **(f)** Same as (a) with indicated strains at 37°C. Right graph: Quantification of indicated cells growth at 37°C on glycerol media relative to the *WT* strain. Mean ± SEM from 3 independent experiments. *P<0.05, **P<0.01, ***P<0.001, ****P<0.0001, ns, not significant (Mann-Whitney t-test).

In this context, decreasing Dnm1 expression during respiratory growth may impact the formation of Ringo constrictions and lead to tubular networks. The *DNM1* gene was thus placed under the control of the *ADH1* promoter (Fig. 4c), which is documented to be repressed under respiratory conditions ^30^. As a control, the *CYC1* promoter which is activated upon respiration ^31^ was used to mimic endogenous expression of Dnm1 (Fig.4c). High expression of Dnm1 from the *CYC1* promoter led to formation of Ringo networks in glycerol media (Fig. 4d). However, lower expression from the *ADH1* promoter resulted in a decrease of Ringo networks upon respiratory growth (YPG), from 49% in *CYC1* to 28% in *ADH1*, and a concomitant significant increase of cells with tubular networks, from 10% in *CYC1* to 39% in *ADH1* (Fig. 4e). Respiratory growth remained unaffected at 23°C and 30°C for *ADH1* cells and at all temperatures for *CYC1* cells (Fig. 4f and S8f). However, at 37°C, the decreased formation of Ringo networks upon Dnm1 expression from the *ADH1* promoter induced a 49% respiratory growth delay as compared to *WT* cells (Fig. 4f). These results indicate that affecting formation of the Ringo mitochondrial morphology by decreasing Dnm1 expression promotes partial respiratory growth inhibition. Thus, the Ringo morphology is not essential, but is required for optimal respiration, which supports a causal link between mitochondrial form and function.

### Involvement of the Ringo morphology in maintenance of mitochondrial DNA homeostasis during respiration

To further evaluate the mechanistic link between the Ringo phenotype and optimal oxidative phosphorylation, we assessed the expression of distinct subunits of OXPHOS complexes upon the formation of Ringo and Muskaan mitochondrial networks. We monitored the expression of Cox1, Cox2 and Cox4, three distinct OXPHOS subunits of complex IV, and Por1, an outer membrane mitochondrial porin in *WT* and *dnm1Δ* cells upon fermentative and respiratory conditions (Fig. 5 a-c). We observed that while expression of Por1 and Cox4 was unaffected by *DNM1* ablation (Fig. 5a), the levels of Cox1 (Fig. 5b) and Cox2 (Fig. 5c) were significantly decreased in respiratory conditions. The disruption of complex IV stoichiometry may be the first indication that Ringo networks are required for optimal oxidative phosphorylation. Yet, a more intriguing observation is that Cox1 and Cox2 are encoded by mitochondrial DNA (mtDNA) whereas Cox4 and Por1 are encoded by the nuclear genome. This suggests that the Ringo mitochondrial morphology may participate in mtDNA homeostasis during respiration.

**Fig. 5.**
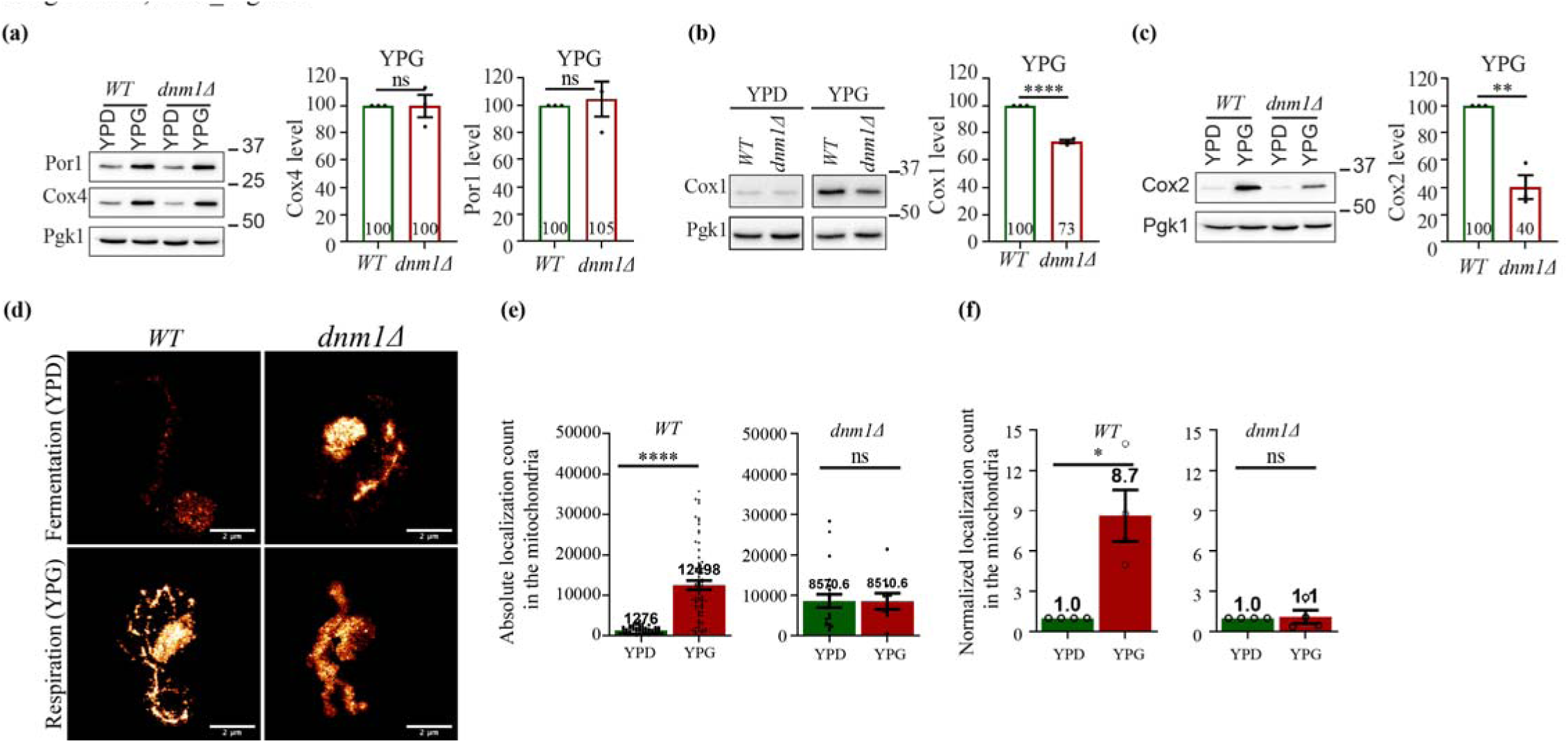
The Ringo morphology regulates OXPHOS components expression through maintenance of mtDNA homeostasis. **(a-c)** Total protein extracts prepared from *WT* and *dnm1*Δ cells in YPD or YPG analyzed by immunoblotting as indicated. MW (kDa) on the right. Right graphs: Cox4, Por1, Cox1 and Cox2 levels normalized to Pgk1 in *dnm1*Δ (red) relative to the *WT* strains (green) in YPG. Mean ± SEM from 3 independent experiments. *P<0.05, **P<0.01, ***P<0.001, ****P<0.0001, ns, not significant (Unpaired t-test). **(d)** Mitochondrial-DNA staining using Hoechst-PAINT in *WT* or *dnm1*Δ cells during fermentation and respiration. Scale bar, 2 μm. **(e)** Absolute count of Hoechst mitochondrial localizations in *WT* and *dnm1*Δ cells (>22 cells) during fermentation (YPD, green) and respiration (YPG, red). **(f)** Same as (e) comparing the amounts of localizations in YPG relative to YPD. In (E) and (F), Mean ± SEM from 3 independent experiments. *P<0.05, **P<0.01, ***P<0.001, ****P<0.0001, ns, not significant (Mann-Whitney t-test).

To address this possibility, we visualized mtDNA using Hoechst-based point accumulation in nanoscale topography (PAINT), a super-resolution technique suited to imaging DNA ^32^. We visualized intracellular DNA by Hoechst-PAINT during fermentative and respiratory growth in wild-type and *dnm1Δ* cells (Fig. 5d). As expected from the known increase in mtDNA copy number upon respiratory as compared to fermentative growth ^33^, Hoechst-PAINT showed that mtDNA localizations significantly increase upon respiration (Fig. 5e and 5f; WT). This mtDNA increase is unlikely caused by additional mito-ER contacts and mitochondrial fission events ^34,35^ since both occur at equivalent frequencies in fermentation and respiration conditions (Fig. 3d and S7a). Consequently, if the Ringo morphology participates in the increase of mtDNA during respiration, this increase should be blocked by abrogation of Dnm1 expression. Hoechst-PAINT revealed that in absence of Dnm1, high amounts of mtDNA localizations are present in fermentation, upon hyperfused mitochondrial networks formation (Fig. 5e; *dnm1Δ*). However, the increase in localizations seen in *WT* cells during respiration with Ringo networks (Fig. 5e and 5f; WT) was totally abolished in *dnm1Δ* cells with Muskaan mitochondria (Fig. 5e and 5f; *dnm1Δ*). These data are consistent with a role of the Ringo morphology in regulating the amount of mtDNA during respiration.

## Discussion

The correlation of mitochondrial morphology and respiration in fungi has been the subject of an ongoing debate ^5,6,8,9^. However, the limitations of confocal microscopy in characterizing mitochondrial morphology at high-resolution have thus far hindered a clear understanding of the impact of mitochondrial form on oxidative phosphorylation ^5,6^. Our results reveal that regular constrictions mediated by Dnm1 along the mitochondrial network during respiration give rise to the formation of a previously unknown morphology we refer to as “Ringo”. The data support a model in which the Ringo morphology maintains optimal levels of mtDNA to favor stoichiometric assembly of OXPHOS complexes. The mechanism of mitochondrial constrictions is distinct from mitochondrial division, since recruitment of Dnm1 to outer membranes occurs independent of contacts with the ER. The signals that trigger Dnm1 targeting to mitochondria in the absence of ER contacts remain elusive. Yet, the Mdv1 adaptor together with increased expression of Dnm1 during respiration likely plays a role in this process. Intriguingly, a similar increase of the mammalian dynamin DRP1 accompanied with extensive remodelling of mitochondrial morphology has been observed in skeletal muscle during exercise ^36^. This suggests that mechanistic and functional features of the Ringo mitochondrial morphology may be evolutionally conserved, serving to optimize oxidative phosphorylation across species.

## Methods

### Yeast culture and transformation

The *saccharomyces cerevisiae* strains used in this study are listed in Supplementary Table 1. Yeast cells were grown and transformed using standard method ^37^. In the indicated strains, *DNM1*, *MMM1*, *OM45*, *TOM70* and other genes were chromosomally deleted or C-terminally tagged using conventional homologous recombination approaches ^38,39^. Where indicated, strains were transformed with plasmids listed in Supplementary Table 2 under selection of interest. Precultures were launched overnight at 30°C in YP complete or Synthetic minimal media completed with 2% Dextrose. According to the experimental conditions, growth media was switched with distinct carbon sources at 2% Dextrose, Glycerol, Ethanol, Galactose or Raffinose. Cells were diluted to achieve at least 3 cell divisions and reach the OD600 = 0.6 to 0.8 per mL of culture at the time of image acquisition.

For culture media switch experiments from Extended Data Fig. 2c-d, cells pre-cultured in YPD/YPG media were diluted into YP media containing distinct dextrose concentrations (0.2%, 0.5%, and 2%) at OD600 = 0.6 per mL of culture. Cells were then allowed to grow before image acquisition at various time points.

For respiration inhibition experiments from Extended Data Fig. 8a, cells were cultured in YPD/YPG media at 30°C to mid-log phase (OD600 = 0.6-0.7 per mL of culture). AntimycinA (Sigma, A8674) reconstituted in ethanol was then added to the culture at 1 µM concentration. For control samples, equal volumes of ethanol were added to cultures.

### Western blot preparation and quantification

Cells were collected during the exponential growth phase and total protein extracts were prepared by the NaOH and trichloroacetic acid (TCA) lysis method ^40^. Proteins were separated by SDS-PAGE and transferred onto Membrane Amersham protran 0,45 µm (Amersham, 10600002). The primary antibodies used for immunoblotting were monoclonal anti-Pgk1 (1/20,000, AbCam, ab113687), monoclonal anti-GFP (1/1,000, Roche, 11814460001), polyclonal anti-Dnm1 (1/1,000, generated by Covalab), monoclonal anti-Por1 (1/10,000, Invitrogen, 459500), anti-Cox1 (1/1,000, AbCam, ab110270), anti-Cox2 (1/1,000, AbCam, ab110271) and anti-Cox4 (1/1,000, AbCam, ab110272). The primary antibodies were detected by incubation with horseradish peroxidase (HRP)-conjugated anti-mouse or anti-rabbit secondary antibodies (Sigma-Aldrich, A9044 and A5278), followed by incubation with the Clarity Western ECL Substrate (Bio-Rad, 1705060). Immunoblotting images were acquired with a Gel Doc XR+ (Bio-Rad) and quantified with the Image Lab 6.0 software (Bio-Rad). The cytosolic protein Pgk1 was used as a loading control to normalize loading of other proteins relative to the WT conditions.

### Image and statistical analysis

For image analysis, we used ImageJ ^41^ and wrote dedicated macro to do the analysis. This Macro was used for all the sample in the same experiment. Results are shown as mean ± standard error of the mean (SEM). Graphs were created and statistical analysis was performed using Graphpad Prism (v8.0). The specific statistical tests are indicated in the figure legends. P value corresponds to *P<0.05, **P<0.01, ***P<0.001, ****P<0.0001, ns, not significant.

### Sample preparation for live cell imaging

Yeast cells were harvested upon reaching the exponential phase (OD600 = 0.6-0.8 per mL of culture). 1 mL of cell culture was centrifuged and resuspended in 30 µL of fresh media. 8 µL of cell suspension was gently placed on 25 mm glass coverslip (#CG15XH, Thorlabs, Germany). Cells were immobilized using a small agar pad containing culture media (*e.g.*, YPD, YPG, SD, SG etc. depending on the experimental condition) which was placed gently on the cell suspension. This resulted in a monolayer of immobilized yeast cells optimal for live cell imaging. The coverslip was ultimately placed in the 25 mm magnetic chamber (#CM-B25-1, Chamlide magnetic chambers) for imaging.

### Spinning disc confocal

Confocal cross-sectional images (Fig. 1a) were acquired using Spinning-disk UltraView VOX (Perkin Elmer) system equipped with a confocal scanning head (CSU X1; Yokogawa), a 100x 1.4 NA oil immersion objective, and EMCCD cameras (ImageEM C9100; Hamamatsu

Photonics) controlled by Volocity software. The GFP and mcherry channel images were acquired using 488 and 561 lasers with 10% and 25% of laser power, respectively, and a maximum of 300 ms of exposure time. The time-lapse movies were recorded for 3 minutes at 10-second intervals. Post-processing of images was performed using ImageJ (National Institute of Health) open-source software.

### Structured illumination microscopy

Cells were imaged using commercial Zeiss LSM 780 Elyra microscope (Carl Zeiss, Germany) controlled by the Zen software. The microscope was equipped with an oil immersion 100x Plan-Apochromat objective with a 1.46 numerical aperture and an additional 1.6x lens. For detection, an EMCCD Andor Ixon 887 1K camera was used. One SIM image was reconstructed from nine images acquired from three different phases and three different angles. Acquisition parameters were adapted to optimize the signal to noise ratio according to yeast strains and protein under investigation. For z-stack acquisition, we kept the z-interval between 80-150 nm depending on the strain and experimental requirement. SIM images were reconstructed with ZEN software and then channel alignment was performed using 100 nm TetraSpeck fluorescent beads (Cat#T7279, Invitrogen) embedded in the same conditions as the sample.

### Mitochondrial morphology quantification

In Figures 1a-b, 2a-c, 4b, 4d-e, S1c, S2b-e, S8a and S8d-e, we quantified the morphology visually and considered only those cells in which more than 70% of mitochondrial network was visible. A z-stack of images was converted into z-projection image using Maximum intensity projection. Each field contained two channels, one for OM (Tom70-GFP) and one for matrix (mt-mcherry). We mostly focused on GFP labelling of OM. Tubular morphology was identified by two parallel threads forming oval shaped close to cortex of cell. Ringo morphology was identified by series of rings connected throughout the cortex of cells, which appear like Tom70 localized in spiral fashion around the matrix. Ringo morphology was far denser and had branched networks. Fragmented mitochondria were easily identified by separated rings, localized mostly at the center of a yeast cell. Hyperfused mitochondria appeared much thicker than tubular, and partially covered the cells unlike tubular or Ringo mitochondria. It appeared significantly straight structure and localized at the edges of cell. Finally, Muskaan morphology was easily identified by few uneven size rings, much thicker than Ringo mitochondria. Muskaan morphology partially covered the cells like Hyperfused mitochondria. From each field we calculated cells with different morphology and compared as percentage of total in a population of cells.

### Photo Activated Localization Microscopy (PALM)

For PALM imaging, (Fig. 1c and S1b), we used an inverted Nikon Ti-E Eclipse microscope equipped with a 100× 1.49 NA objective (Nikon) with the Perfect Focus System active. A 488-nm wavelength laser with 200 mW power (at 1%) was used to acquire low resolution images before activation using blue light source. Blue LED light source emitting at 405 nm wavelength was used to the photo activation of mEOS. A 561-nm wavelength laser with 500 mW (at 75%) was used to acquire 50,000 images without interval with an exposure time of 50 ms. Images were acquired on an EMCCD camera with 1.5x additional magnification, resulting in pixel size of 106 nm and a field of view (FoV) of 54 μm x 54 μm. EM gain of the EMCCD was set to 300. A 3.7% formaldehyde (#8.18708, Sigma-Aldrich) fixed yeast cells in which Tom70, an outer mitochondrial membrane protein tagged with mEOS2 was mounted on the coverslip using collagen gel. Briefly, 3.6 mg/ml collagen stock (#354236, Corning) was diluted to final matrix concentrations of 2.2 mg/ml and neutralized with 1 M sodium hydroxide (#28-3010, Sigma-Aldrich). Cells were mixed into this collagen solution and a drop was mounted on the top of the coverslips and allowed to polymerize at RT for 30 minutes. Collagen gel was used to immobilize yeast cells because agar pad introduced significant background and non-specific blinking events.

### Single Molecule Localization microscopy (SMLM) image reconstruction

For SMLM data reconstruction (PALM and Hoechst PAINT), the ThunderSTORM ^42^ plugin available in ImageJ ^41^ was used. We chose Wavelet filter (B-Spline) for image filtering, local maximum method for approximate localization of molecule. PSF: Gaussian method was used for sub-pixel localization of molecules with weighted least squares fitting method. We either used fluorescent beads or cross-correlation to perform the drift correction. We filtered out localization that have sigma values below 100 or above 200, or localization that have uncertainty above 50 nm. Localization appearing in consecutive frames separated by less than 60 nm was merged, with the assumption that they are originating from the same molecule. Finally, localization table was saved into “.CSV” format and Normalized Gaussian method was used for image visualization. Final image resolution was in the range of 40-50 nm.

### Transmission Electron microscopy

Yeast cells were processed from adapted OTO protocol ^43^. Briefly, cells were grown in YPD/YPG medium and were fixed by immersion in 2.5% glutaraldehyde, and post-fixed. First in a mix of 1% OsO4 with 1.5% K4Fe(CN)6 (potassium ferrocyanide – reduced osmium) to increase the membranes contrast, then contrast was enhanced by tannic acid 1%. A second post-fixation was performed in 1% OsO4. Subsequently, the samples were dehydrated through increasing graded ethanol series and embedded in epoxy resin (Electron Microscopy Sciences) at 60°C for 48 hrs. The embedded resin block was mounted on a specimen stub using colloidal silver paint (Electron Microscopy Sciences) and sputter coated with 200nm of gold/palladium.

Embedding samples were cut with an ultramicrotome. Ultrathin sections (70 nm) were cut with a Leica Ultracut S microtome and stained with uranyl acetate followed by lead citrate. Contrasted sections were observed using a TEM FEI Tecnai T12 at 120kV.

### TEM images quantification

To generate the graphs in Fig. 1f and 1g from TEM images, we quantified mitochondrial diameter by drawing a line randomly from one end to the other end of mitochondria. The length of the line was then measured. For final plotting, an average of 3 random lines per cell was considered. To count the number of rings and constriction, we visually counted the structure/sites on mitochondria such as those denoted by green and yellow arrows in Fig. 1d and S4. Circularity of mitochondria was set as an important factor for ring characterization. We thus considered not counting oval shaped mitochondria as rings. For constrictions, continuity in the mitochondrial network was the main factor along with the small diameter of mitochondrial tubules.

### Cryo-Electron Tomography

For grids preparation, *S. cerevisiae* cells (*WT* or *dnm1*Δ) were grown in either YPD or YPG media to an OD600 = 0.9-1. 3 µl of cell suspension was applied to a freshly grown discharged holey carbon grid (Quantifoil® #200, R2/2). Grids were immediately blotted from the back side using Whatman filter paper #1 (GE Healthcare) within the automatic plunge freezer EM GP (Leica Microsystems). Grids were stored at liquid nitrogen temperature until further use.

Lamellae were prepared using an Aquilos FIB-SEM system (Thermo Fisher Scientific). Grids were coated with an initial organometallic platinum layer using a gas injection system (GIS) for 10 seconds followed by 20 seconds sputtering coating to add an additional inorganic platinum layer. Samples were tilted to a milling angle of 8o. Milling was done iteratively in 3 steps (1: 1 nA – 5 µm, 2: 0.5 nA - 3µm, 3: 0.3 nA – 1.3 µm) before polishing at 30-50 pA to achieve a final lamella thickness of 100-200 nm. After polishing the EM grids were sputtered again for 2-3 seconds.

For data acquisition, 12 (YPD conditions) and 14 (YPG conditions) tilt series were acquired for the wild type experiments and 12 (YPD conditions) and 16 (YPG conditions) tilt series of the dnm1Δ sample. Tilt series were collected either on a Titan Krios electron microscope (Thermo Fisher Scientific) equipped with a K2 summit electron detector (Gatan) and a Quantum energy filter using a pixel size of 2.682 Å/pixel or, on a Titan Krios G4 electron microscope (Thermo Fisher Scientific) equipped with Selectris X imaging filter and Falcon 4 direct electron detector using a pixel size of 3.037 Å/pixel. Movies were collected in a tilt range of +68 o / -52 o starting at +8 o pretilt with a 2 o tilt increment. Target defocus was set to -2.5 to -4.5 µm. A total electron dose of approximately 120 e/Å2 per tilt series was used following a dose-symmetric tilt scheme in SerialEM ^44,45^.

For tomogram reconstruction and volume segmentation, movies of individual projections were motion-corrected and aligned within SerialEM ^44^. Combined and dose-filtered stacks were aligned and reconstructed using AreTomo ^46^. To improve visualization reconstructed tomograms were binned by 4 and deconvoluted using tom_deconv ^47^. Volume segmentation was done manually within the Amira software package (Thermo Fisher Scientific). Final images were prepared using ChimeraX ^48^.

### Cryo-ET image quantification

Generating the graphs in Fig. 1g and 2e-h from Cryo-ET images was done as follows. To measure the diameter of mitochondria, an average of 3 random lines drawn between both ends of OM was considered for the final value. To quantify the ring and constriction of mitochondria, we drew a line along the OM through all sections of a tomogram. This allowed obtaining 2-dimention projections of tomograms which were used to perform the quantification of rings and constriction as described in the TEM image quantification section above. Tomograms also allowed counting the number of cristae. As a first step, a given tomogram was scanned from bottom to top to appreciate the appearing cristae visually. To measure the length of the cristae, we drew a line through the cristae along the tomogram and took the average to 3 longest cristae for the final value. To quantify the cristae crossing the mitochondria, we scanned the tomogram and counted the cristae which intersect the entire mitochondrial tubule. To quantify the presence of ER at constriction site, we counted those constriction site in which ER was present in close proximity (<75 nm) per tomogram. To perform comparison with SIM images, we took the projection of four SIM images plane (≈300 nm, equivalent to Cryo-ET images) and cropped the same dimension for field of view. To measure the diameter of mitochondria, we drew 3 lines across mitochondria tubules, plotted the line profile and measured the distance between the two peaks. Counting of rings and constriction were done as described in the TEM image quantification section above.

### Fluorescent puncta quantification

GFP puncta quantification of Dnm1 and Mmm1 used in Figures 3a and 3d were achieved on a single plane of SIM images using ImageJ Macro. Initially, a plane from the stack where major portions of mitochondrial networks were detected was selected. The background in both channels was then subtracted. This was done by delimiting a region of empty space in the Field of View (FoV) to measure the mean intensity from both channels in this region and subtract it from respective channels. To quantify the puncta, two duplicate images were created and a Gaussian blur of 5 and 7 pixels (the size of the blur depends on the size of the puncta) was introduced. After subtracting the two images, a threshold was applied manually to generate a binary image. Mitochondria were subsequently quantified by using Gaussian blur followed by manual thresholding. We selected the individual yeast cells in the FoV and created a Region of interest (ROI) around each cell, making sure that each ROI contained a single cell and avoiding overlap between two ROIs. The number of puncta in each ROI was counted to finally represent the number of puncta per cell. In parallel, the puncta density on mitochondria was measured. For this purpose, the area of mitochondria in each ROI was evaluated to obtain the ratio of puncta count relative to the area of mitochondria. All the parameters were kept constant for each condition while analyzing the images. This pipeline allowed performing a semi-automated quantification.

### Mitochondrial fission quantification

Mitochondrial fission events quantification in Fig. 3e, 3f and S7a was performed as follows. We generated a strain in which Dnm1 is tagged with mcherry and Mmm1 with GFP (MCY2199) in which mitochondria were labelled with Mitotracker (#M22426, Invitrogen). This setup allowed following and quantifying the presence of Dnm1-mCherry and Mmm1-GFP at sites of mitochondrial fission. This quantification was performed on raw time-lapse images that were acquired for 3-minute duration at every 10 second time interval. Before analysis, the image contrast was adjusted in all three channels but in settings where the mitochondria channel was the only one visible. After identifying a fission event, Dnm1 and Mmm1 were checked for their presence in the proximity at the fission site. The fission events were classified into 4 categories including fission event where (1) both Dnm1 and Mmm1 (2) only Dnm1, (3) only Mmm1, and (4) none are present. The results were plotted as the percentage of total for each category. To generate graphs from Fig. 3g, the localization of Dnm1-GFP puncta on the mitochondrial tubules was visually quantified using single planes from z-stacks. After adjusting the image contrast, puncta were categorized into two groups, one localized randomly on mitochondrial tubule (similar to Fig. 3g top image) and another one where puncta specifically localized on mitochondrial constrictions (similar to Fig. 3h bottom image). We plotted the graph as percentage of total of puncta localized in each group.

### PreCOX4 intensity measurement

The analysis from Extended Data Fig. 7b and 8b was performed using an ImageJ Macro on single planes from SIM images with the MCY1607 and MCY2161 strains where Tom70 is tagged with GFP and Precox4 with mcherry. Planes from z-stacks where major portions of mitochondrial networks was detected were selected. The background was subtracted as described in the puncta quantification section above. The Tom70-GFP channel was employed to segment mitochondrial networks because of the strong GFP signal. Gaussian blur followed by manual thresholding was used to create a binary image of mitochondria. ROI was created to segment the single cell, considering that ROIs were not overlapping each other. Mean intensity of precox4-mCherry in the mitochondrial ROI was measured. Ultimately, MS Excel and GraphPad prism were used to assemble the data and generate the plots. Each dot in the plots from Extended Data Fig. 7b and 8b corresponds to the mean intensity of Precox4-mCherry in one cell. Bars and error bars represent the mean value and SEM, respectively.

### Spot assay preparation and quantification

Cultures grown overnight in minimal synthetic medium with Dextrose (SD) were pelleted, resuspended at OD600=1 per mL of culture, and serially diluted (1:10) three times in water. Five microliters of the dilutions were spotted on SD or minimal synthetic medium with Glycerol (SG) plates and grown for 2 to 4 days (Dextrose) or 3 to 6 days (Glycerol) at 23, 30 or 37°C. “Microarray Profile” plugin for ImageJ was used to quantify each spot in Fig. 4f. Data reported are the mean and s.d. (error bars) from three independent experiments.

### Hoechst PAINT

Hoechst-PAINT acquisition was performed on yeast spheroplasts. To prepare spheroplasts, cells were grown at mid log phase (0.6-0.8 OD600). 1 mL of 3.7% formaldehyde was added to the 10 mL of cell culture and incubated for 20 minutes at 300C. Cells were then washed with PBS twice, then re-suspended into 3 mL of spheroplast buffer containing 50 mM β-mercaptoethanol (#63689, Sigma-Aldrich) + Zymolyase (#ZE1005, Zymo Research; Orange, CA) at 3 µL/10 OD of culture in PBS. Re-suspended cells were incubated for 30 minutes at 30°C in a 15 mL tube placed horizontally in a rotating incubator. After two washes with PBS cells were permeabilized using 0.5% of Triton™ X-100 (#T8787, Sigma-Aldrich) in PBS for 30 minutes on a shaker.

Finally, spheroplasts were washed 3 times with PBS and stored in 300 μl of PBS at 40C. Spheroplasts were stable for 2 weeks for further processing and imaging. To immobilize spheroplasts, 25 mm coverslip was coated with 1 mg/ml Concanavalin (#C5275, Sigma-Aldrich) in PBS for 2 hours, then washed 3 times with PBS, dried overnight and stored at 4°C in PBS. 30 μL of spheroplasts suspension was mixed with 270 μL of PBS and spread over 25 mm coverslip for 1 hour and then washed 3 times with PBS. The coverslip was placed into Chamlide magnetic chambers and 200 pM of JF646-Hoechst, novel fluorogenic DNA stain (Gift from Janelia farm) conjugated with Janelia Fluor® 646 dye in 1 mL of PBS was added to the chamber. Image acquisition was performed on the same system used for PALM imaging. A 647-nm wavelength laser with 500 mW (at 60 %) was used to acquire 100,000 images without interval with an exposure time of 100 ms. EM gain of the EMCCD was set to 300.

### Hoechst-PAINT data quantification

To quantify information from PAINT data in Fig. 5e-f, the .CSV file generated by ThunderSTORM was used and displayed as a Histogram. Even though we used a strain in which Matrix is tagged with mCherry (MCY1949), the mCherry signal was not strong enough to allow automatic segmentation of mitochondria. Consequently, the ROI corresponding to mitochondria in each cell was manually drawn, to measure the area and number of localizations from mitochondrial DNA, which allowed calculating the absolute and density of localizations in each cell.

## Data availability

All data are available in the main text or the supplementary materials.

## Acknowledgments

Research in the Cohen laboratory is supported by the Agence Nationale de la Recherche (ANR) grants, labex DYNAMO (ANR-11-LABX-0011-DYNAMO), MOMIT (ANR-17-CE13-0026-01) and MITOFUSION (ANR-19-CE11-0018). C.Z. also acknowledges funding by ANR (ANR-17-CE13-0026-01, ANR-21-CE45-003-02 and ANR-16-CONV-0005) and by Institut Pasteur. We gratefully acknowledge the Imagopole—Citech of Institut Pasteur (Paris, France) as well as the France–BioImaging infrastructure network supported by the French National Research Agency (ANR-10–INSB–04; Investments for the Future) for the use of the Zeiss LSM 780 Elyra PS1 microscope. We thank the EM facility of the Max Planck Institute of Biophysics (Frankfurt, Germany), in particular Werner Kühlbrandt for critical reading of the manuscript and Sonja Welsch, Oezkan Yildiz and Juan Castillo for computing support. We also thank the Max Planck Society and Deutsche Forschungsgesellschaft (FOR2848). We are grateful for support for electron microscopy equipment from the French Government Programme Investissements d’Avenir France BioImaging (FBI, N° ANR-10-INSB-04-01) and the French gouvernement (Agence Nationale de la Recherche) Investissement d’Avenir programme, Laboratoire d’Excellence “Integrative Biology of Emerging Infectious Diseases” (ANR-10-LABX-62-IBEID). We thank Nadège Cayet for her help to section embedded samples and TEM acquisition.

## Author contributions

MKS acquired and analyzed all SIM, PALM and PAINT data. LC and NBT prepared all plasmids and strains and performed all western blots and serial dilution assays. ML brought assistance to MKS in PALM and PAINT data acquisition. PB and AM acquired all TEM data. LD and CK acquired all Cryo-ET data. MKS analyzed all TEM and Cryo-ET data. MKS and MMC conceptualized the project. MMC and CZ supervised MKS. MMC and MKS wrote the manuscript with inputs from CZ and LD and all authors reviewed and edited the paper.

## Competing interests

Authors declare that they have no competing interests.

---

**Extended Data Fig. 1:**
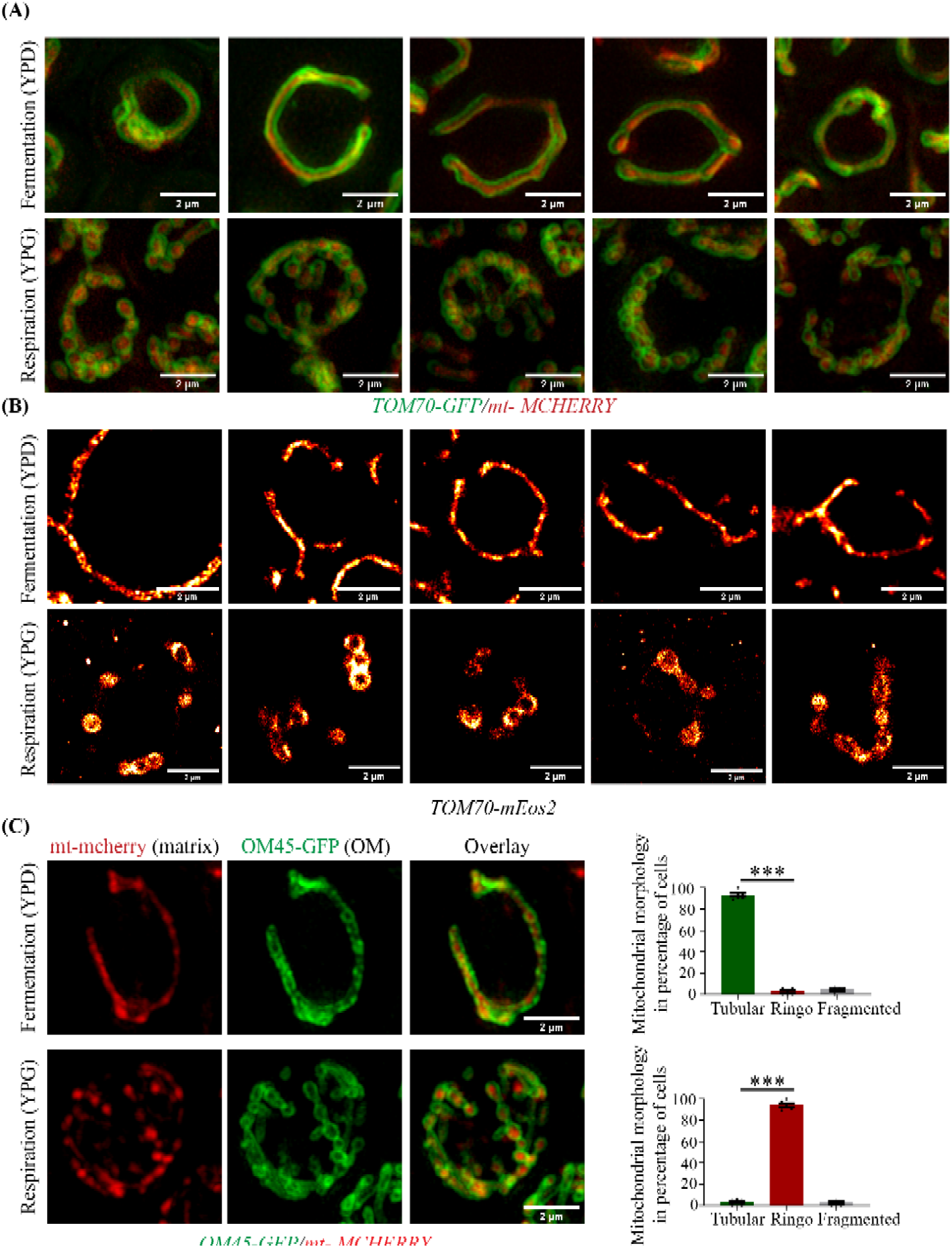
SIM and PALM detection of the Ringo mitochondrial morphology does not depend on the protein used to label OMs. **(a)** Five distinct SIM acquisitions of cells labeled for mitochondrial matrix (mt-mCherry) and Outer Membranes (Tom70-GFP) in fermentation (top) or respiration (bottom). Scale bar, 2 µm . **(b)** Five distinct PALM images of cells labeled with TOM70-mEOS2 in fermentation (top) or respiration (bottom). Scale bar, 2 µm. **(c)** SIM acquisitions of cells labeled for mitochondrial matrix (mt-mCherry) and Outer Membranes (OM45-GFP) in fermentation (top) or respiration (bottom). Scale bar, 2 µm. Right graphs: percentage of cells with Tubular, Ringo or Fragmented mitochondria. Mean ± SEM from >177 cells in 3 independent experiments. *P<0.05, **P<0.01, ***P<0.001, ****P<0.0001, ns, not significant (Mann-Whitney t-test). Note that the respiratory Ringo morphology forms independent of the outer membrane protein used to label mitochondrial outer membranes (*i.e.* OM45 or Tom70).

**Extended Data Fig. 2:**
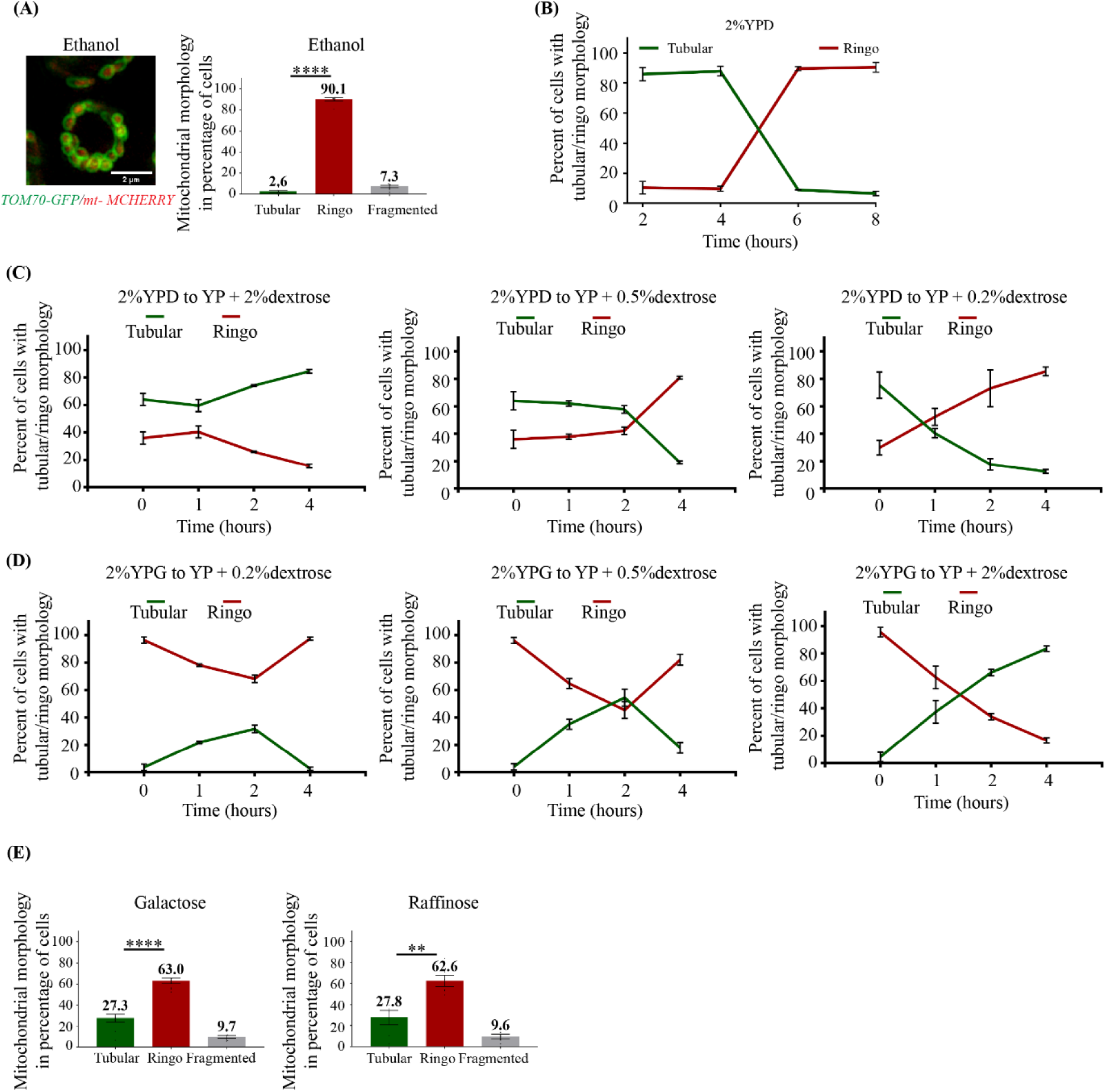
The Ringo morphology dominates in all respiratory conditions. **(a)** SIM acquisitions of WT cells labeled for mitochondrial matrix (mt-mCherry) and Outer Membranes (Tom70-GFP) in Ethanol media. Scale bar, 2 µm. Right graph: percentage of cells with Tubular, Ringo or Fragmented mitochondria. Mean ± SEM from >703 cells in 3 independent experiments. ****P<0.0001, (Mann-Whitney t-test). Note that the Ringo morphology dominates in the Ethanol respiratory media. **(b)** Percentage of WT cells labeled with mt-mCherry and Tom70-GFP with Tubular or Ringo mitochondria during 8 hours culture in 2% YPD. Mean ± SEM from >78 cells in 1 experiment. Note that mitochondrial morphology progressively switches from Tubular to Ringo after 4 hours of culture which is presumably caused by progressive dextrose depletion. **(c)** Percentage over 4 hours of WT cells (labeled with mt-mCherry and Tom70-GFP) with Tubular or Ringo mitochondria. Cells were pre-cultured in 2% YPD (fermentative) and switched to YP media with various dextrose concentration (2%, 0.5%, and 0.2%). Mean ± SEM from >221 cells in 3 independent experiments. Note that the lower the glucose concentration, the faster the switch from Tubular to Ringo morphology occurs. **(d)** Same as (c) but cells were pre-cultured in 2% YPG (respiration) and switched to YP media containing various concentration glucose such as, 0.2%, 0.5%, and 2%. Mean ± SEM from >145 cells in 3 independent experiments. Note that the higher the glucose concentration, the faster the switch from Ringo to Tubular morphology takes place. **(e)** Percentage of WT cells (labeled with mt-mCherry and Tom70-GFP) with Tubular, Ringo or Fragmented mitochondria in Galactose (left) and Raffinose (right) media. Mean ± SEM from >374 cells in 3 independent experiments.

**Extended Data Fig. 3:**
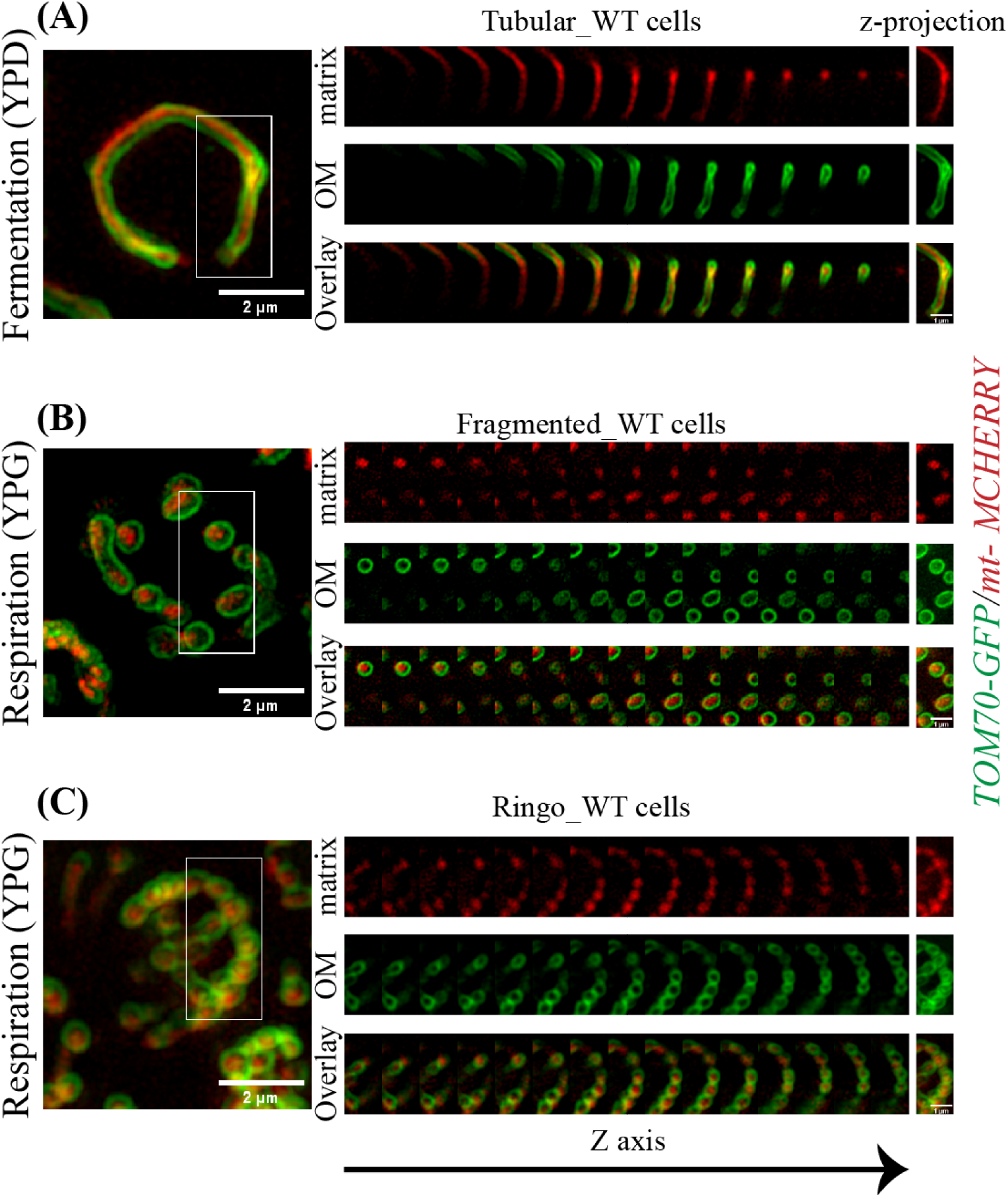
Matrix within Ringo mitochondria is continuous. Maximum intensity projection (MIP) of z-stack SIM acquisitions from cells labeled for mitochondrial matrix (mt-mCherry) and Outer Membranes (Tom70-GFP) with distinct mitochondrial morphologies. Series represents top-to-bottom individual slices of the stack along Z-axis. MIP of inset is shown at the end of each row. Scale bar, 2 µm and for inset, 1 µm. **(a)** Tubular morphology during fermentation. The matrix in red in continuous. **(b)** Fragmented morphology during respiration. The matrix in red is not continuous. **(c)** Ringo morphology during respiration. The matrix in red seems continuous.

**Extended Data Fig. 4:**
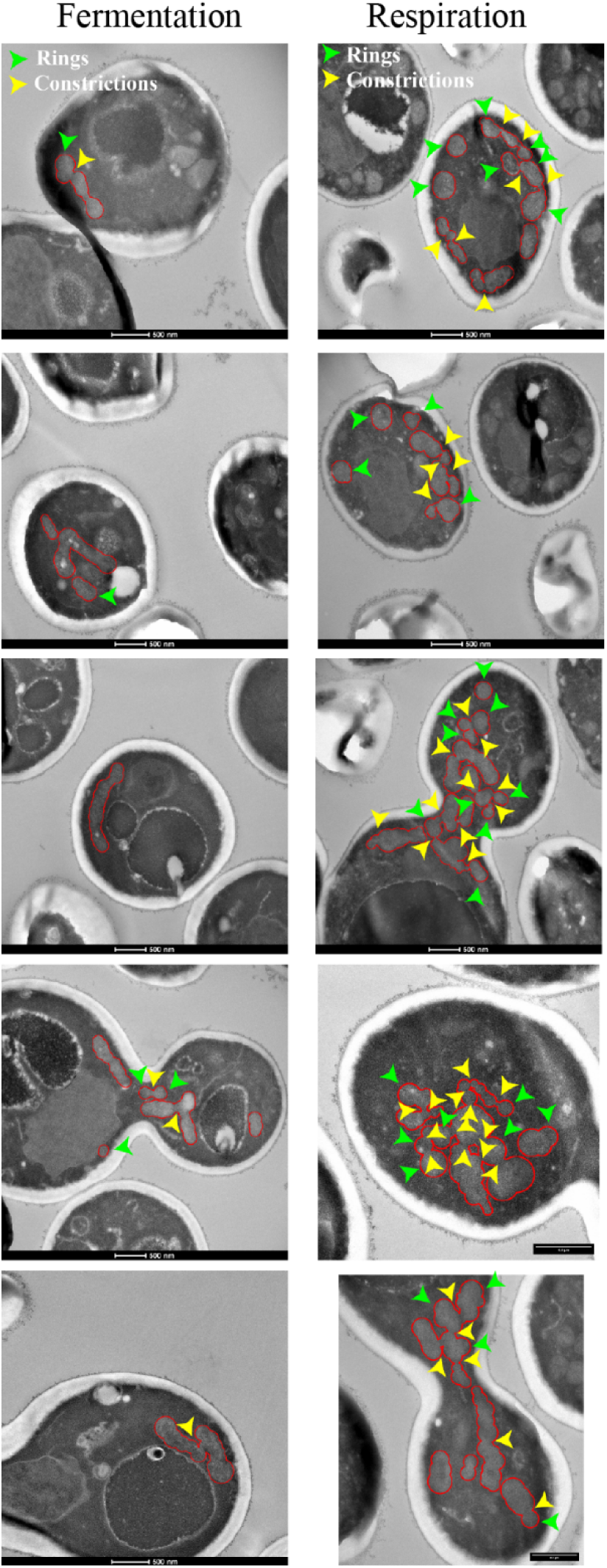
Transmission Electron Microscopy (TEM) reveals ultrastructural differences between Tubular and Ringo mitochondrial networks. Various examples of TEM micrographs of cells in fermentation (Scale bar, 1 µm) or respiration (Scale bar, 1 µm or 500 nm). Mitochondria are delimited by red demarcations. Rings and constrictions are indicated by green or yellow arrowheads, respectively. Note the strong increase of mitochondrial constrictions within Ringo mitochondrial networks.

**Extended Data Fig. 5:**
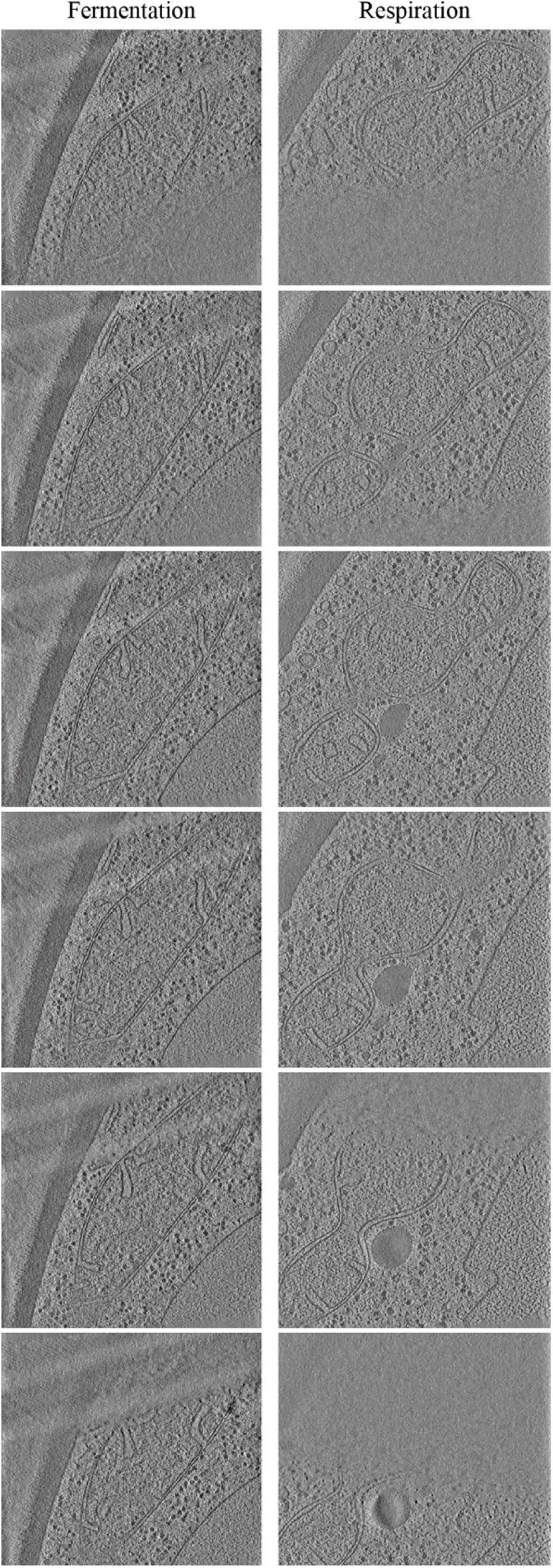
Cryo-Electron Tomography (Cryo-ET) confirms ultrastructural differences between Tubular and Ringo mitochondrial networks. Top to bottom slices through tomographic volumes of cell in fermentation (left) and respiration (right). Scale bars, 500 nm.

**Extended Data Fig. 6:**
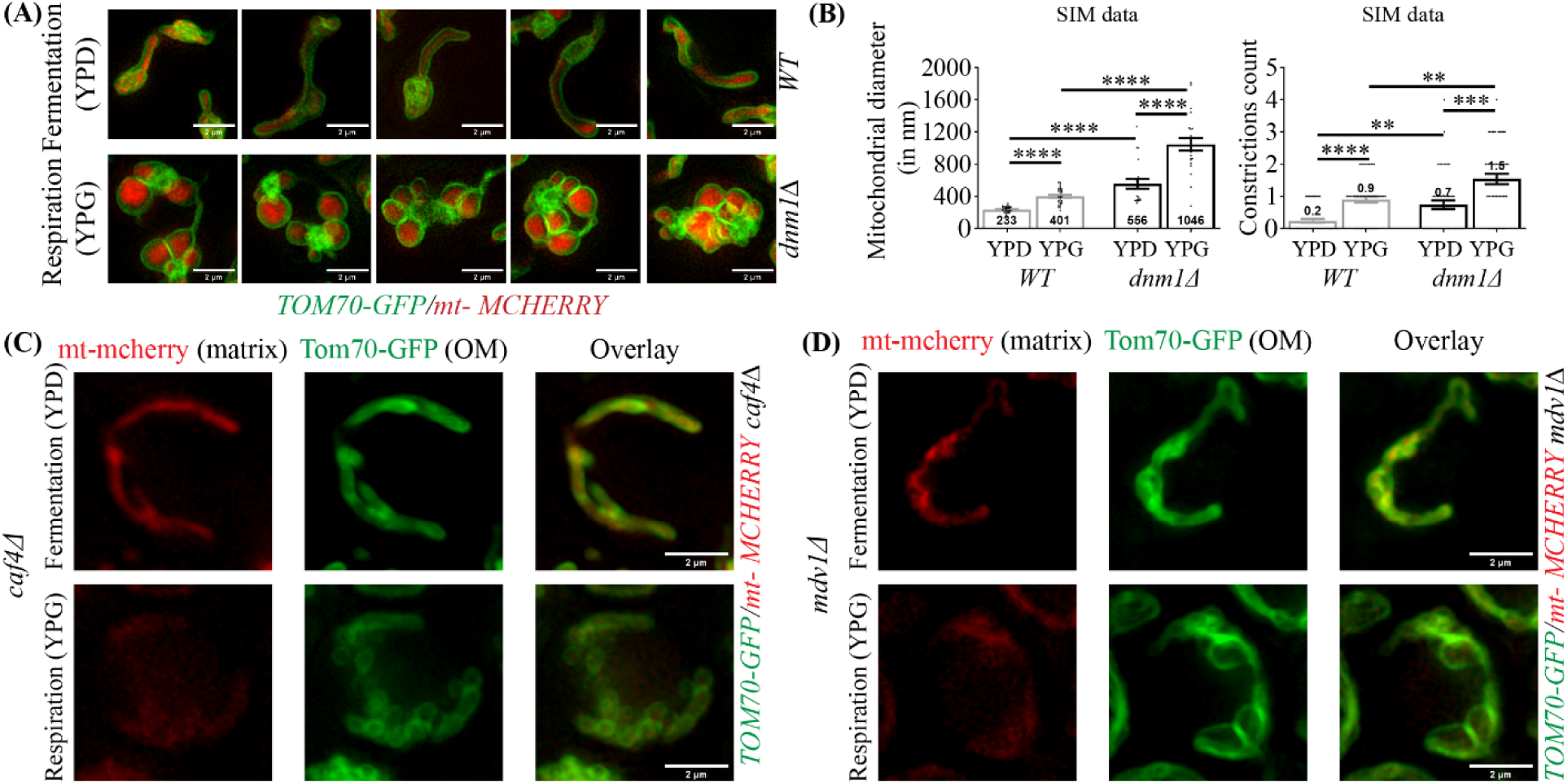
The Muskaan mitochondrial morphology is generated in the absence of either Dnm1 or Mdv1. **(a)** Five distinct SIM acquisitions of *WT* or *dnm1*Δ cells labeled for mitochondrial matrix (mt-mCherry) and Outer Membranes (Tom70-GFP) in fermentation (top) or respiration (bottom). Scale bar, 2 µm. **(b)** Average mitochondrial diameter (left), and constriction counts per cell (right) as quantified in SIM acquisitions from *WT* or *dnm1*Δ cells in YPD or YPG. Mean ± SEM from >25 cells in 3 independent experiments. *P<0.05, **P<0.01, ***P<0.001, ****P<0.0001, (Mann-Whitney t-test). Note that mitochondria with the Muskaan morphology are distinct from mitochondria with Ringo morphology in terms and diameter and amount of apparent constrictions. **(c and d)** SIM acquisitions of *caf4*Δ (b) or *mdv1*Δ (c) cells labeled for mitochondrial matrix (mt-mCherry) and Outer Membranes (Tom70-GFP) in fermentation (top) or respiration (bottom). Scale bar, 2 µm. Note that the Muskaan morphology is seen in the absence of Mdv1 but not in the absence of Caf4 where Ringo networks dominate.

**Extended Data Fig. 7:**
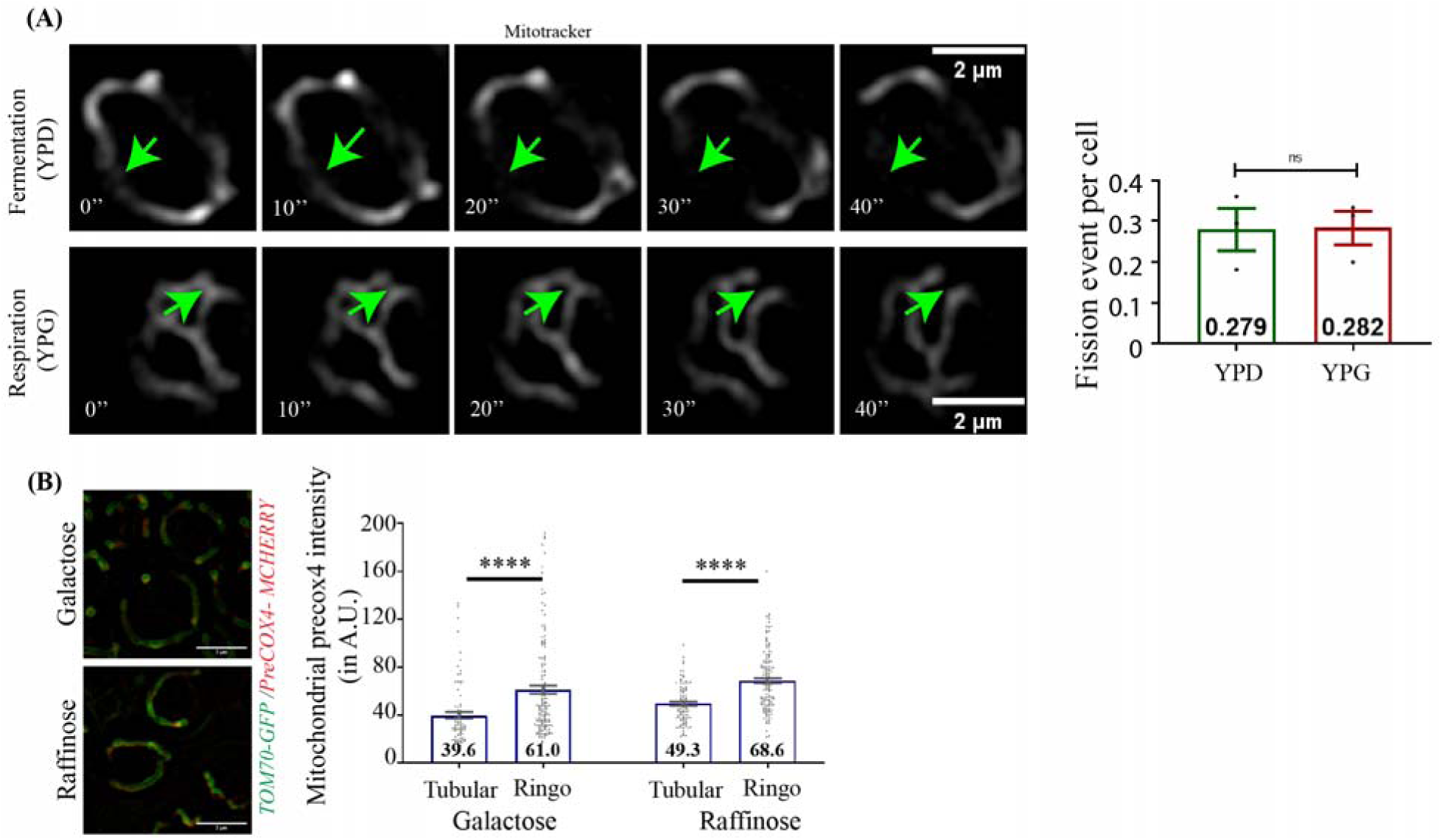
Mitochondrial fission efficiency in dextrose or glycerol media and mitochondrial import in galactose or raffinose media. **(a)** Screenshot of SIM Time-lapse from Dnm1-mcherry, Mmm1-GFP, mitotracker (grey) triple-labeled cells during fermentation and respiration. Green arrows indicate fission events. Quantitation of fission events per cell as mean ± SEM from >175 cells and 3 independent experiments. ns, not significant (Mann-Whitney t-test). Note that Fission efficiency is equivalent in fermentation and respiration. **(b)** SIM acquisitions of cells labeled for the mitochondrial import marker precox4-mcherry and Outer Membranes (Tom70-GFP) in Galactose and Raffinose media. Scale bar, 3 µm. Right graph: Mitochondrial precox-4 mcherry intensity (in Absolute Units) within Tubular and Ringo mitochondria in galactose and raffinose media. Mean ± SEM from >227 cells in 3 independent experiments. ***P<0.001, ****P<0.0001, (Mann-Whitney t-test). Note that Ringo mitochondria that respire show increased transport of precox-4 mcherry as compared to tubular mitochondria that do not respire.

**Extended Data Fig. 8: Related to Figure 4.**
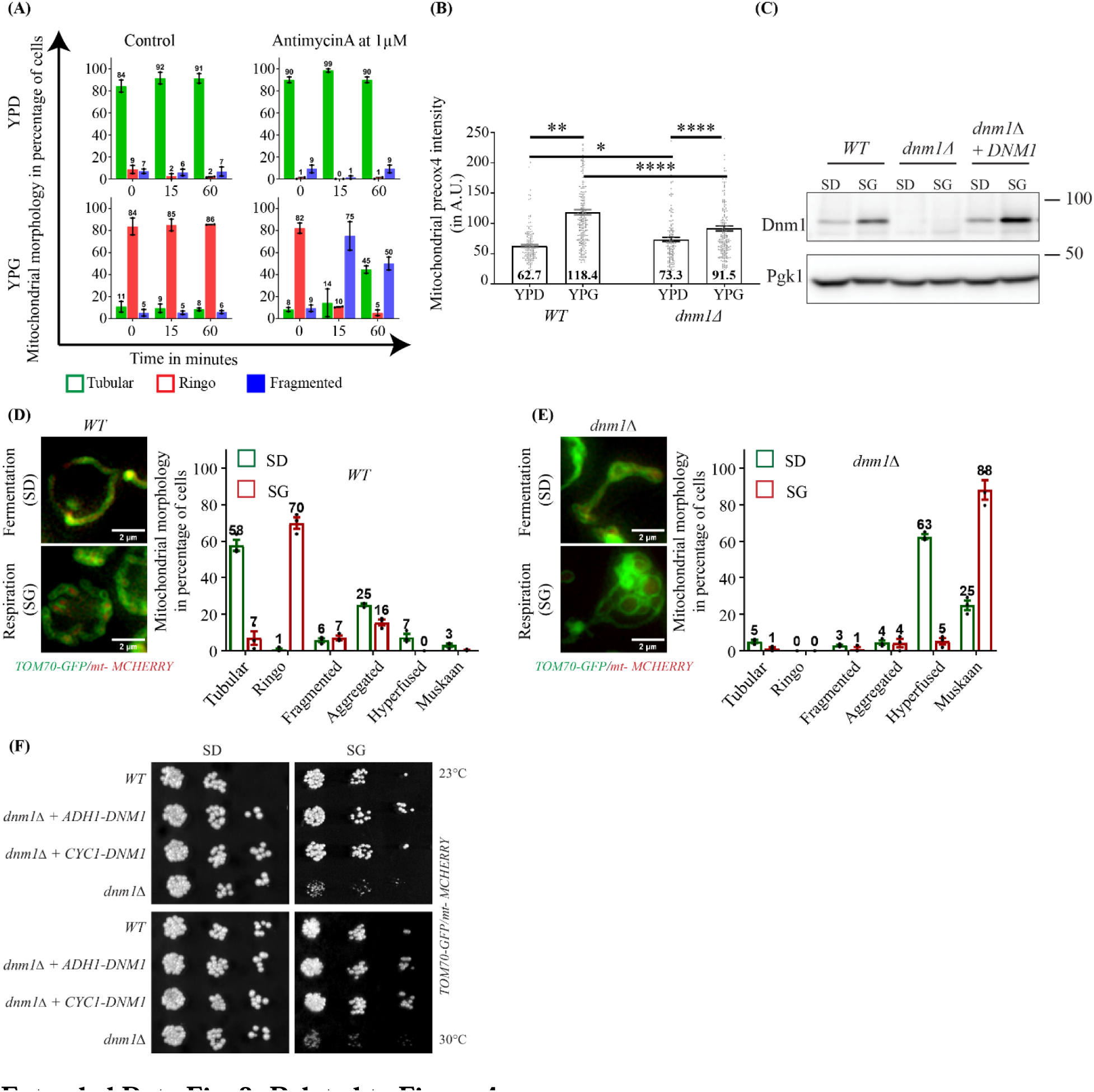
**(a)** Percentage of cells with Tubular, Ringo or Fragmented mitochondria in control or AntimycinA (1µM) treated cells during fermentation and respiration. Mean ± SEM from >156 cells in 3 independent experiments. Note that AntimycinA treatment induces fragmentation of Ringo networks. **(b)** Mitochondrial precox-4 mcherry intensity (in Absolute Units) within WT and dnm1Δ cells labelled with Tom70-GFP in fermentation and respiration. Mean ± SEM from >156 cells in 3 independent experiments. *P<0.05, **P<0.01, ****P<0.0001, (Mann-Whitney t-test). Note that Ringo mitochondria show increased transport of precox-4 mcherry as compared to Muskaan mitochondria. **(c)** Total protein extracts prepared from in WT, dnm1Δ or dnm1Δ+DNM1 cells in Dextrose (SD) or Glycerol (SG) media and analyzed by immunoblotting as indicated with anti-Dnm1 and anti-Pgk1. MW (kDa) indicated on the right. **(d)** SIM acquisitions of WT cells labeled for mitochondrial matrix (mt-mCherry) and Outer Membranes (Tom70-GFP) in fermentation or respiration. Scale bar, 2 µm. Right: Percentage of cells with Tubular, Ringo, Fragmented, Aggregated, Hyperfused or Muskaan mitochondria. Mean ± SEM from >180 cells in 3 independent experiments. **(e)** Same as (d) with dnm1Δ cells. **(f)** Dextrose and Glycerol serial dilutions of WT, dnm1Δ+ADH1-DNM1, dnm1Δ+CYC1-DNM1, and dnm1Δ strains at 23, and 30°C.

**Supplementary Table 1.** Strains Used in the Study

| Background and number | Yeast Name | Genotype | Occurrence in the study | Reference |
| --- | --- | --- | --- | --- |
| W303 (MCY553) | WT | Mat a, *ura3-1 trp1-1 leu2-3,112 his3-11,15 can1-100 RAD5 ADE2* | 1d-h, ED4, ED5 |  |
| W303 (MCY1949) | *TOM70-GFP/mt- MCHERRY* | Mat a, *TOM70-GFP::CaURA3 ura3-1 trp1-1, mt-mcherry::LEU2,112 his3-11,15 can1-100 RAD5 ADE2* | 1a, 1b, 2a, 4a, 5a-f, S1a, S2, ED3, ED6, ED7b, ED8a-f | This study |
| W303 (MCY1497) | *TOM70-mEos2* | Mat a, *ura3-1 trp1-1 leu2-3,112 his3-11,15 can1-100 RAD5 ADE2 TOM70::mEos2-KanMX6* | 1c, ED1b | This study |
| W303 (MCY1940) | *OM45-GFP/mt- MCHERRY* | Mat a, *OM45-GFP::KanMX6 ura3-1 trp1-1 mt-MCHERRY::LEU2,112 his3-11,15 can1-100 RAD5 ADE2* | ED1c | This study |
| W303 (MCY1940) | *TOM70-GFP /mt- MCHERRY/ dnm1Δ* | Mat a, *TOM70-GFP::CaURA3 ura3-1 DNM1::TRP1, mt- MCHERRY::LEU2,112 his3-11,15 can1-100 RAD5 ADE2* | 2a, 2d-h, 3b, 5a-f, ED6a-b | This study |
| W303 (MCY2099) | *TOM70-GFP/mt- MCHERRY caf4Δ* | Mat a, *CAF4::NatMX4 TOM70-GFP::CaURA3 ura3-1 trp1-1, mt- MCHERRY::LEU2,112 his3-11,15 can1-100 RAD5 ADE2* | 2c, ED6c | This study |
| W303 (MCY2097) | *TOM70-GFP::mt- MCHERRY mdv1Δ* | Mat a, *MDV1::NatMX4 TOM70-GFP::CaURA3 ura3-1 trp1-1, mt- MCHERRY::LEU2,112 his3-11,15 can1-100 RAD5 ADE2* | 2d, ED6d | This study |
| W303 (NBT1343) | *TOM70- MCHERRY /DNM1- GFP* | *TOM70:: MCHERRY -Nat DNM1::GFP-TRP ura3-1 trp1-1 leu2-3,112 his3-11,15 can1-100 RAD5 ADE2* | 3a, 3d, 3g | This study |
| W303 (NBT344) | *TOM70- MCHERRY/MMM1- GFP* | *TOM70:: MCHERRY -Nat MMM1::GFP-TRP ura3-1 trp1-1 leu2-3,112 his3-11,15 can1-100 RAD5 ADE2* | 3a, 3b, 3d, 3g | This study |
| W303 (MCY2199) | *Dnm1- MCHERRY /MMM1- GFP* | Mat a, *ura3-1 trp1-1 leu2-3,112 his3-11,15 can1-100 RAD5 ADE2; MMM1-GFP::TRP1 DNM1-MCHERRY::NATNT2* | 3e-f, ED7a | This study |
| W303 (MCY1607) | *TOM70-GFP /PreCOX4- MCHERRY* | Mat a, *TOM70-GFP::CaURA3 PreCox4- MCHERRY::HphMX ura3-1 trp1-1 leu2-3,112 his3-11,15 can1-100 RAD5 ADE2* | ED7b, ED8b | Gift from Zhou XU |
| W303 (MCY2161) | *TOM70-GFP/PreCOX4- MCHERRY/dnm1Δ* | Mat a, *TOM70-GFP::CaURA3 PreCox4- MCHERRY::HphMX ura3-1 DNM1::TRP1 leu2-3,112 his3-11,15 can1-100 RAD5 ADE2* | ED8b | This study |
| W303 (MCY2166) | *TOM70-GFP/mt- MCHERRY* | Mat a, *ura3-1 trp1-1 leu2-3,112 his3-11,15 can1-100 RAD5 ADE2; mt-MCHERRY::LEU2; TOM70-GFP::CaURA3; pRS413* | 4a, 4f, ED8c, ED8d, ED8f | This study |
| W303 (MCY2167) | *TOM70-GFP/mt- MCHERRY/ dnm1Δ + CYC1-DNM1* | Mat a, *ura3-1 trp1-1 leu2-3,112 his3-11,15 can1-100 RAD5 ADE2; mt-MCHERRY::LEU2; TOM70-GFP::CaURA3; dnm1::TRP1; p413CYC1-DNM1* | 4c, 4d, 4f, ED8f | This study |
| W303 (MCY2168) | *TOM70-GFP/mt- MCHERRY/dnm1Δ + ADH-DNM1* | Mat a, *ura3-1 trp1-1 leu2-3,112 his3-11,15 can1-100 RAD5 ADE2; MITO-MCHERRY::LEU2; TOM70-GFP::CaURA3; dnm1::TRP1; p413ADH-DNM1* | 4c, 4e, 4f, ED8f | This study |
| W303<br>(MCY2169) | *TOM70-GFP/mt- MCHERRY/*<br>*dnm1Δ + TEF-DNM1* | Mat a, *ura3-1 trp1-1 leu2-3,112 his3-11,15 can1-100*<br>*RAD5 ADE2; MITO-MCHERRY::LEU2; TOM70-*<br>*GFP::CaURA3; dnm1::TRP1; p413TEF-DNM1* | 3c | This study |
| W303<br>(MCY2170) | *TOM70-GFP/mt- MCHERRY/*<br>*dnm1Δ + DNM1* | Mat a, *ura3-1 trp1-1 leu2-3,112 his3-11,15 can1-100*<br>*RAD5 ADE2; MITO-MCHERRY::LEU2; TOM70-*<br>*GFP::CaURA3; dnm1::TRP1; pRS413-DNM1-DNM1* | 4a, 4b, ED8c | This study |
| W303<br>(MCY2171) | *TOM70-GFP/mt- MCHERRY/*<br>*dnm1Δ* | Mat a, *ura3-1 trp1-1 leu2-3,112 his3-11,15 can1-100*<br>*RAD5 ADE2; MITO-MCHERRY::LEU2; TOM70-*<br>*GFP::CaURA3; dnm1::TRP1; pRS413* | 4a, 4f, ED8c,<br>ED8e, ED8f | This study |

**Supplementary Table 2.** Plasmids Used in the Study

| **Name (Collection number)** | **Description** | **Reference** |
| --- | --- | --- |
| pRS413 | CEN, HIS3, Amp | 49 |
| p413CYC1-*DNM1* | CEN, CYC1 promoter-*DNM1*, HIS3, Amp | This study |
| p413ADH-*DNM1* | CEN, ADH promoter-*DNM1*, HIS3, Amp | This study |
| p413TEF-*DNM1* | CEN, TEF promoter-*DNM1*, HIS3, Amp | This study |
| pRS413-*DNM1-DNM1* | CEN, *DNM1* promoter-*DNM1*, HIS3, Amp | This study |

## Notes

### Competing Interest Statement

The authors have declared no competing interest.

## References

1. Quintana-Cabrera, R. & Scorrano, L. Determinants and outcomes of mitochondrial dynamics. Mol Cell 83, 857–876 (2023).

2. Tilokani, L., Nagashima, S., Paupe, V. & Prudent, J. Mitochondrial dynamics: overview of molecular mechanisms. Essays Biochem 62, 341–360 (2018).

3. Ozeir, M. & Cohen, M. M. From dynamin related proteins structures and oligomers to membrane fusion mediated by mitofusins. Biochim Biophys Acta Bioenerg 1863, 148913 (2022).

4. Sesaki, H. & Jensen, R. E. Division versus fusion: Dnm1p and Fzo1p antagonistically regulate mitochondrial shape. J Cell Biol 147, 699–706 (1999).

5. Visser, W. et al. Effects of growth conditions on mitochondrial morphology in Saccharomyces cerevisiae. Antonie Van Leeuwenhoek 67, 243–253 (1995).

6. Egner, A., Jakobs, S. & Hell, S. W. Fast 100-nm resolution three-dimensional microscope reveals structural plasticity of mitochondria in live yeast. Proc Natl Acad Sci U S A 99, 3370–3375 (2002).

7. Bennett, C. F., Latorre-Muro, P. & Puigserver, P. Mechanisms of mitochondrial respiratory adaptation. Nat Rev Mol Cell Biol 23, 817–835 (2022).

8. Zheng, F. et al. Glucose starvation induces mitochondrial fragmentation depending on the dynamin GTPase Dnm1/Drp1 in fission yeast. J Biol Chem 294, 17725–17734 (2019).

9. Church, C. & Poyton, R. O. Neither respiration nor cytochrome c oxidase affects mitochondrial morphology in Saccharomyces cerevisiae. J Exp Biol 201, 1729–1737 (1998).

10. Friedman, J. R. et al. ER tubules mark sites of mitochondrial division. Science 334, 358–62 (2011).

11. Kornmann, B. et al. An ER-mitochondria tethering complex revealed by a synthetic biology screen. Science 325, 477–81 (2009).

12. Bleazard, W. et al. The dynamin-related GTPase Dnm1 regulates mitochondrial fission in yeast. Nat Cell Biol 1, 298–304 (1999).

13. Smirnova, E., Griparic, L., Shurland, D. L. & van der Bliek, A. M. Dynamin-related protein Drp1 is required for mitochondrial division in mammalian cells. Mol Biol Cell 12, 2245–56 (2001).

14. Zhang, P. & Hinshaw, J. E. Three-dimensional reconstruction of dynamin in the constricted state. Nat Cell Biol 3, 922–6 (2001).

15. Ingerman, E. et al. Dnm1 forms spirals that are structurally tailored to fit mitochondria. J Cell Biol 170, 1021–7 (2005).

16. Rolland, F., Winderickx, J. & Thevelein, J. M. Glucose-sensing and -signalling mechanisms in yeast. FEMS Yeast Res 2, 183–201 (2002).

17. Gancedo, J. M. Yeast carbon catabolite repression. Microbiol Mol Biol Rev 62, 334–361 (1998).

18. Söhngen, N. L. & Coolhaas, C. THE FERMENTATION OF GALACTOSE BY SACCHAROMYCES CEREVISIAE. J Bacteriol 9, 131–141 (1924).

19. Goffrini, P., Ferrero, I. & Donnini, C. Respiration-dependent utilization of sugars in yeasts: a determinant role for sugar transporters. J Bacteriol 184, 427–432 (2002).

20. Guaragnella, N. et al. Yeast growth in raffinose results in resistance to acetic-acid induced programmed cell death mostly due to the activation of the mitochondrial retrograde pathway. Biochim Biophys Acta 1833, 2765–2774 (2013).

21. Cerveny, K. L., McCaffery, J. M. & Jensen, R. E. Division of mitochondria requires a novel DMN1-interacting protein, Net2p. Mol Biol Cell 12, 309–21 (2001).

22. Fekkes, P., Shepard, K. A. & Yaffe, M. P. Gag3p, an outer membrane protein required for fission of mitochondrial tubules. J Cell Biol 151, 333–40 (2000).

23. Hoppins, S., Lackner, L. & Nunnari, J. The machines that divide and fuse mitochondria. Annu Rev Biochem 76, 751–80 (2007).

24. Tieu, Q., Okreglak, V., Naylor, K. & Nunnari, J. The WD repeat protein, Mdv1p, functions as a molecular adaptor by interacting with Dnm1p and Fis1p during mitochondrial fission. J Cell Biol 158, 445–52 (2002).

25. Cerveny, K. L. & Jensen, R. E. The WD-repeats of Net2p interact with Dnm1p and Fis1p to regulate division of mitochondria. Mol Biol Cell 14, 4126–39 (2003).

26. Griffin, E. E., Graumann, J. & Chan, D. C. The WD40 protein Caf4p is a component of the mitochondrial fission machinery and recruits Dnm1p to mitochondria. J Cell Biol 170, 237– 48 (2005).

27. Needs, H. I. et al. Interplay between Mitochondrial Protein Import and Respiratory Complexes Assembly in Neuronal Health and Degeneration. Life (Basel) 11, 432 (2021).

28. Vowinckel, J., Hartl, J., Butler, R. & Ralser, M. MitoLoc: A method for the simultaneous quantification of mitochondrial network morphology and membrane potential in single cells. Mitochondrion 24, 77–86 (2015).

29. Starkov, A. A. & Fiskum, G. Myxothiazol induces H(2)O(2) production from mitochondrial respiratory chain. Biochem Biophys Res Commun 281, 645–650 (2001).

30. Denis, C. L., Ferguson, J. & Young, E. T. mRNA levels for the fermentative alcohol dehydrogenase of Saccharomyces cerevisiae decrease upon growth on a nonfermentable carbon source. J Biol Chem 258, 1165–1171 (1983).

31. Hörtner, H. et al. Regulation of synthesis of catalases and iso-1-cytochrome c in Saccharomyces cerevisiae by glucose, oxygen and heme. Eur J Biochem 128, 179–184 (1982).

32. Spahn, C. K. et al. A toolbox for multiplexed super-resolution imaging of the E. coli nucleoid and membrane using novel PAINT labels. Sci Rep 8, 14768 (2018).

33. Galeota-Sprung, B., Fernandez, A. & Sniegowski, P. Changes to the mtDNA copy number during yeast culture growth. R Soc Open Sci 9, 211842 (2022).

34. Murley, A. et al. ER-associated mitochondrial division links the distribution of mitochondria and mitochondrial DNA in yeast. Elife 2, e00422 (2013).

35. Lewis, S. C., Uchiyama, L. F. & Nunnari, J. ER-mitochondria contacts couple mtDNA synthesis with mitochondrial division in human cells. Science 353, aaf5549 (2016).

36. Moore, T. M. et al. The impact of exercise on mitochondrial dynamics and the role of Drp1 in exercise performance and training adaptations in skeletal muscle. Mol Metab 21, 51–67 (2019).

37. Sherman, F., Fink, G. & Hicks, J. Methods in yeast genetics, 1986.

38. Longtine, M. S. et al. Additional modules for versatile and economical PCR-based gene deletion and modification in Saccharomyces cerevisiae. Yeast 14, 953–61 (1998).

39. Gueldener, U., Heinisch, J., Koehler, G. J., Voss, D. & Hegemann, J. H. A second set of loxP marker cassettes for Cre-mediated multiple gene knockouts in budding yeast. Nucleic Acids Res 30, e23 (2002).

40. Volland, C., Urban-Grimal, D., Geraud, G. & Haguenauer-Tsapis, R. Endocytosis and degradation of the yeast uracil permease under adverse conditions. J Biol Chem 269, 9833– 41 (1994).

41. Girish, V. & Vijayalakshmi, A. Affordable image analysis using NIH Image/ImageJ. Indian J Cancer 41, 47 (2004).

42. Ovesný, M., Křížek, P., Borkovec, J., Svindrych, Z. & Hagen, G. M. ThunderSTORM: a comprehensive ImageJ plug-in for PALM and STORM data analysis and super-resolution imaging. Bioinformatics 30, 2389–2390 (2014).

43. Seligman, A. M., Wasserkrug, H. L. & Hanker, J. S. A new staining method (OTO) for enhancing contrast of lipid--containing membranes and droplets in osmium tetroxide--fixed tissue with osmiophilic thiocarbohydrazide(TCH). J Cell Biol 30, 424–432 (1966).

44. Mastronarde, D. N. Automated electron microscope tomography using robust prediction of specimen movements. J Struct Biol 152, 36–51 (2005).

45. Hagen, W. J. H., Wan, W. & Briggs, J. A. G. Implementation of a cryo-electron tomography tilt-scheme optimized for high resolution subtomogram averaging. J Struct Biol 197, 191– 198 (2017).

46. Zheng, S. et al. AreTomo: An integrated software package for automated marker-free, motion-corrected cryo-electron tomographic alignment and reconstruction. J Struct Biol X 6, 100068 (2022).

47. Tegunov, D. & Cramer, P. Real-time cryo-electron microscopy data preprocessing with Warp. Nat Methods 16, 1146–1152 (2019).

48. Pettersen, E. F. et al. UCSF ChimeraX: Structure visualization for researchers, educators, and developers. Protein Sci 30, 70–82 (2021).

49. Sikorski, R. S. & Hieter, P. A system of shuttle vectors and yeast host strains designed for efficient manipulation of DNA in Saccharomyces cerevisiae. Genetics 122, 19–27 (1989).

