## Supplementary figures and images for "A constricted mitochondrial morphology optimizes respiration"

### Extended Data Fig. 1

Singh M. K., et al._Extended Data Figure1

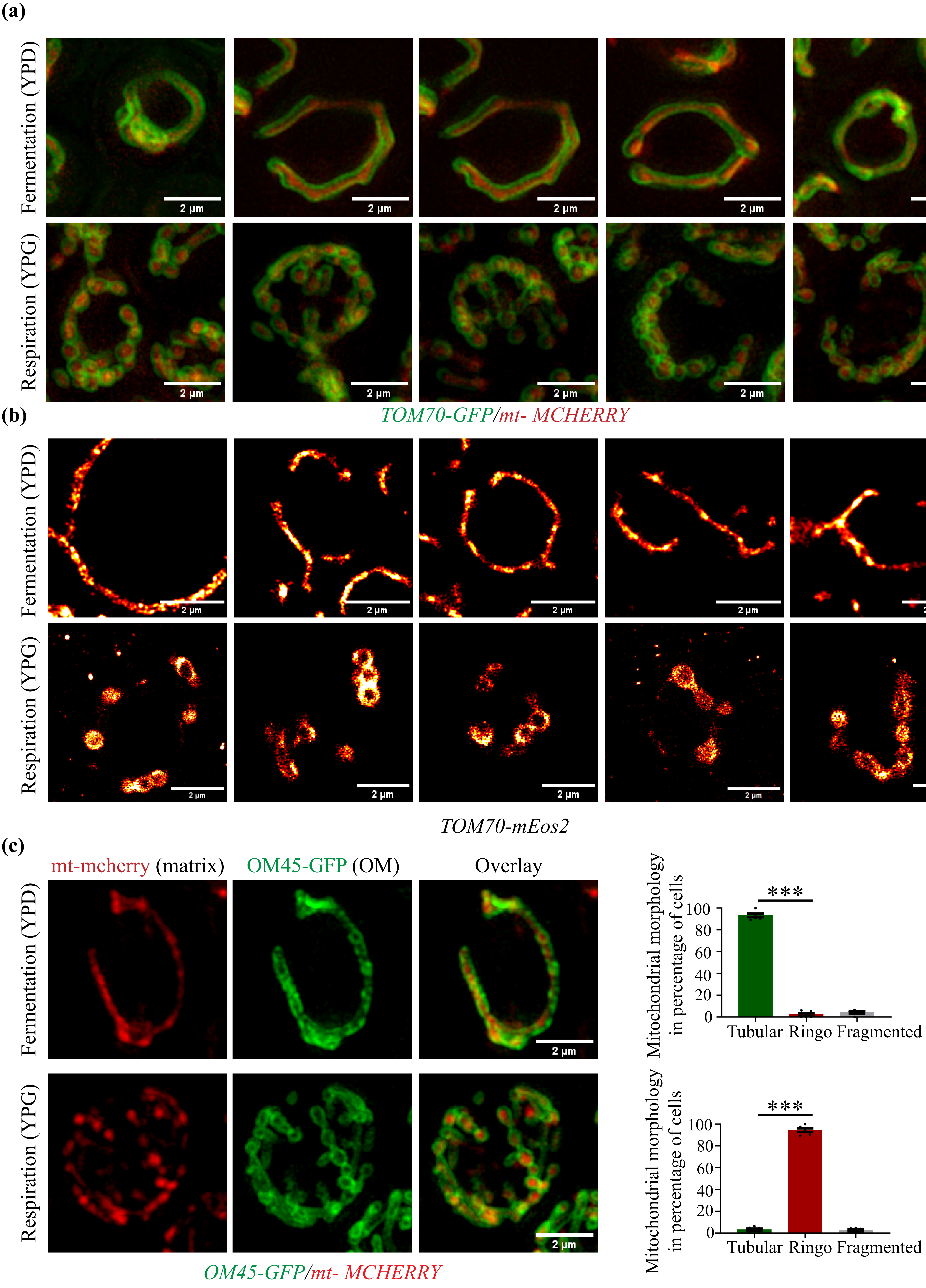

### Extended Data Fig. 2

Singh M. K., et al._Extended Data Figure 2

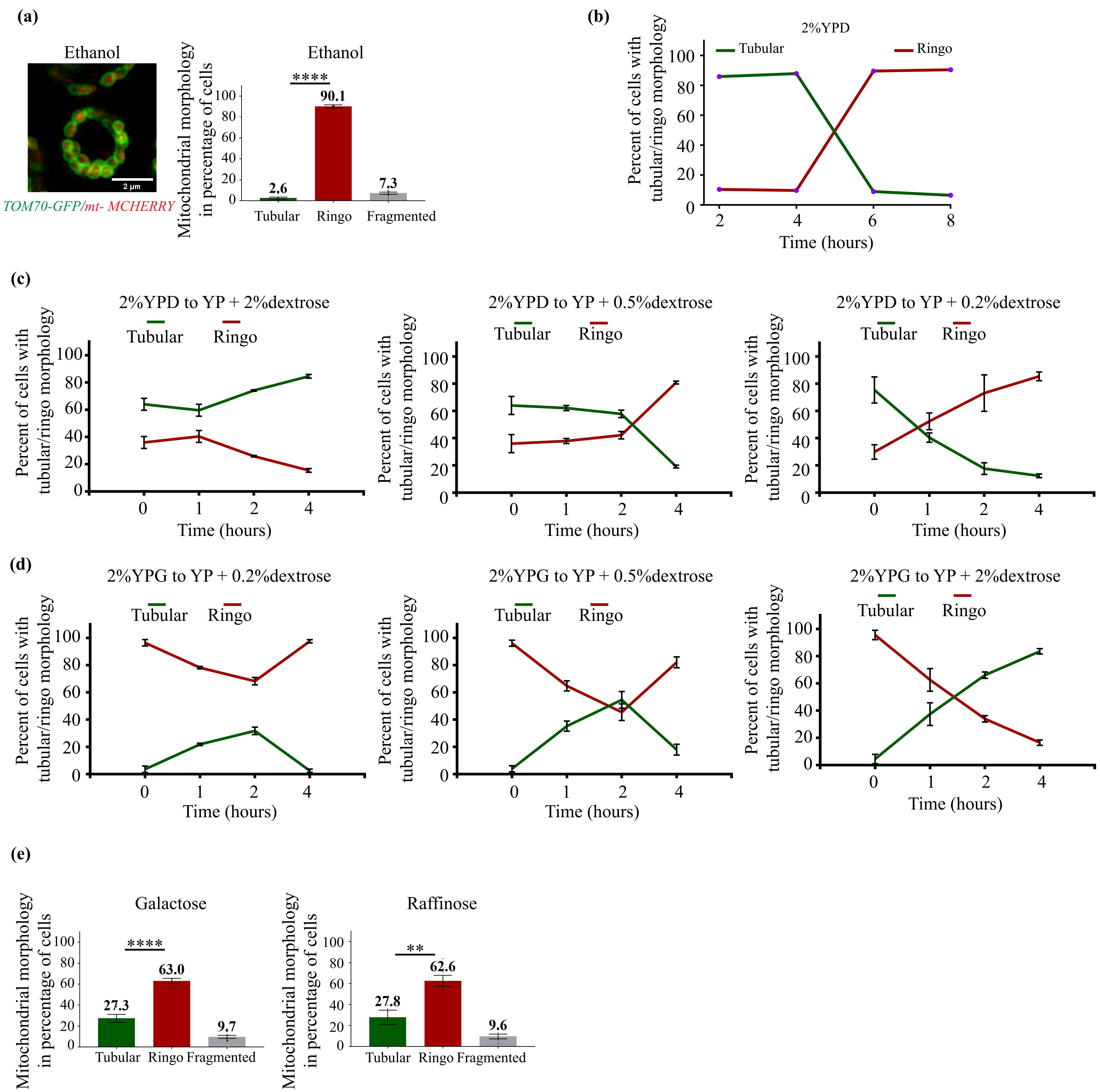

### Extended Data Fig. 3

#### Singh M. K., et al._Extended Data Figure 3

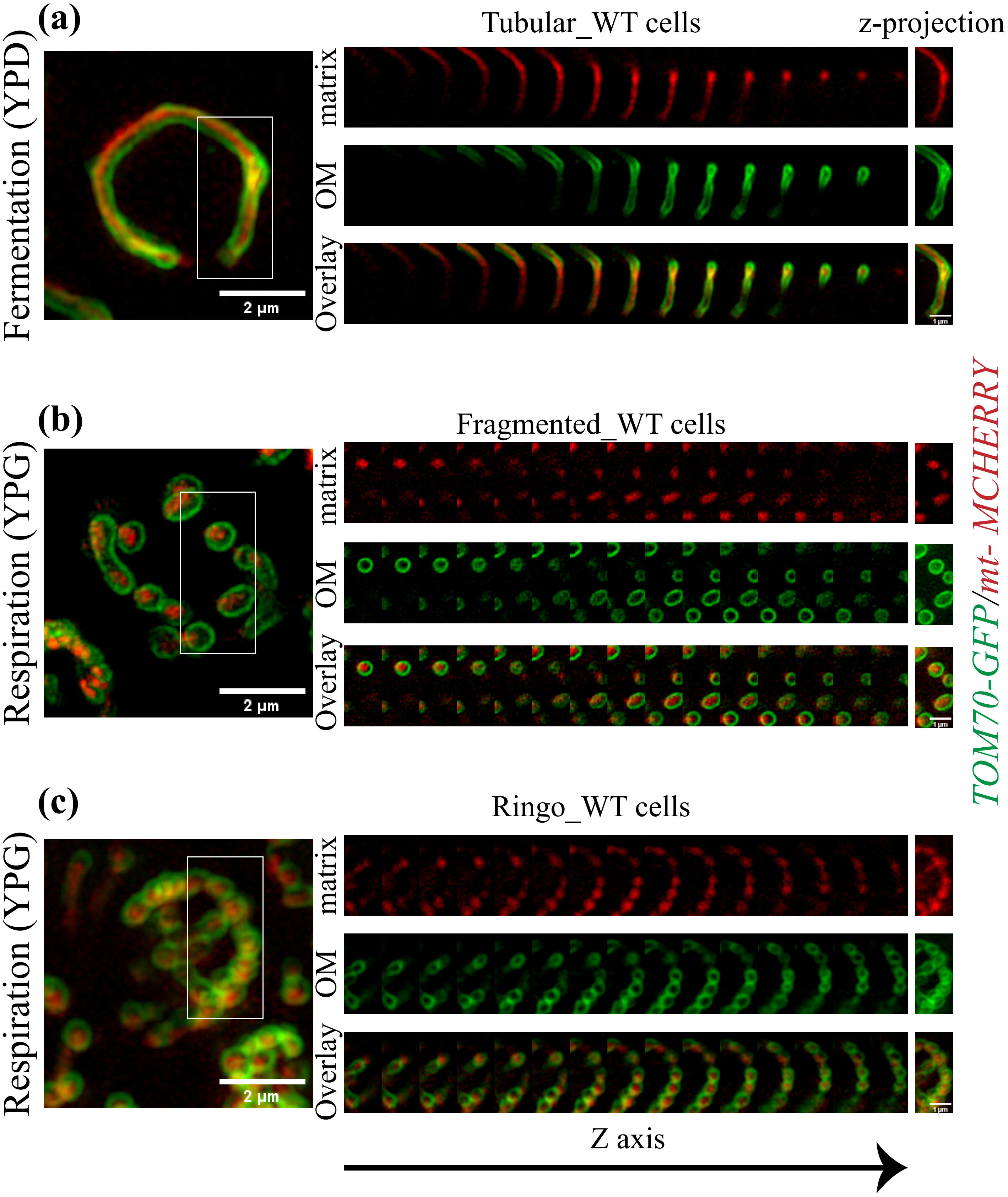

### Extended Data Fig. 4

Singh M. K., et al._Extended Data Figure 4

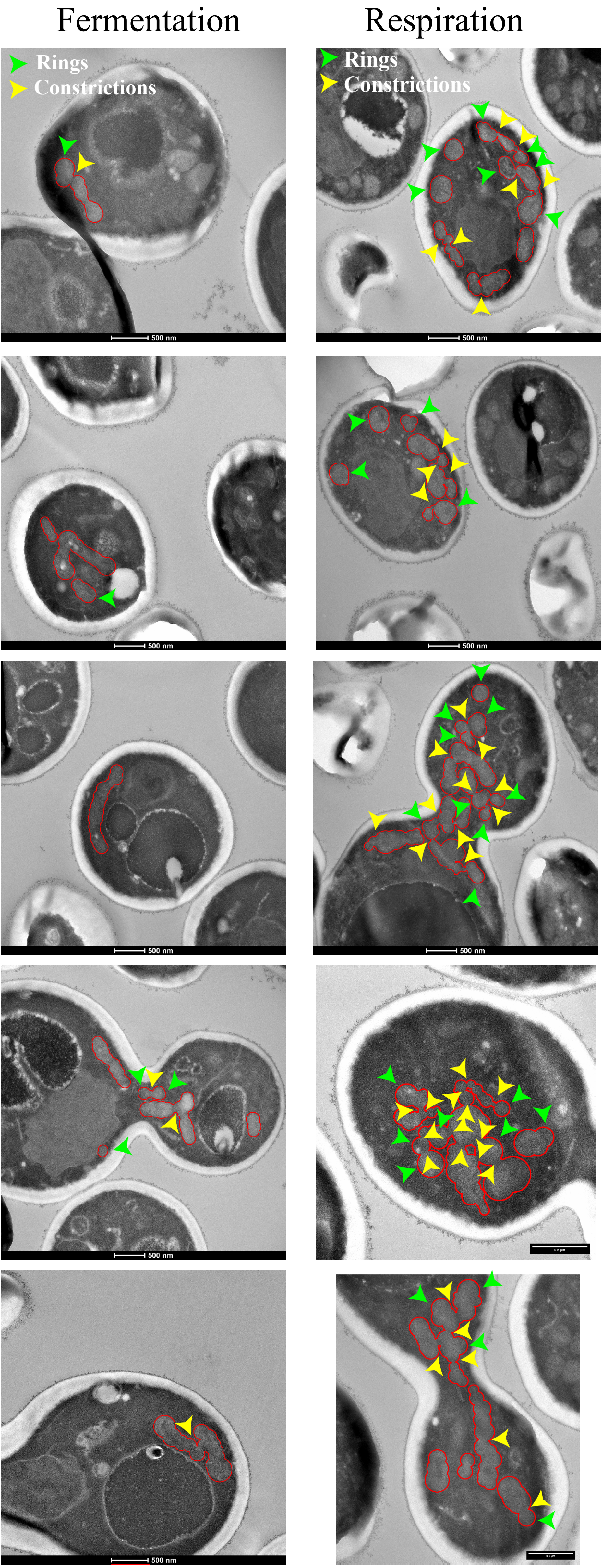

### Extended Data Fig. 5

Singh M. K., et al._Extended Data Figure 5

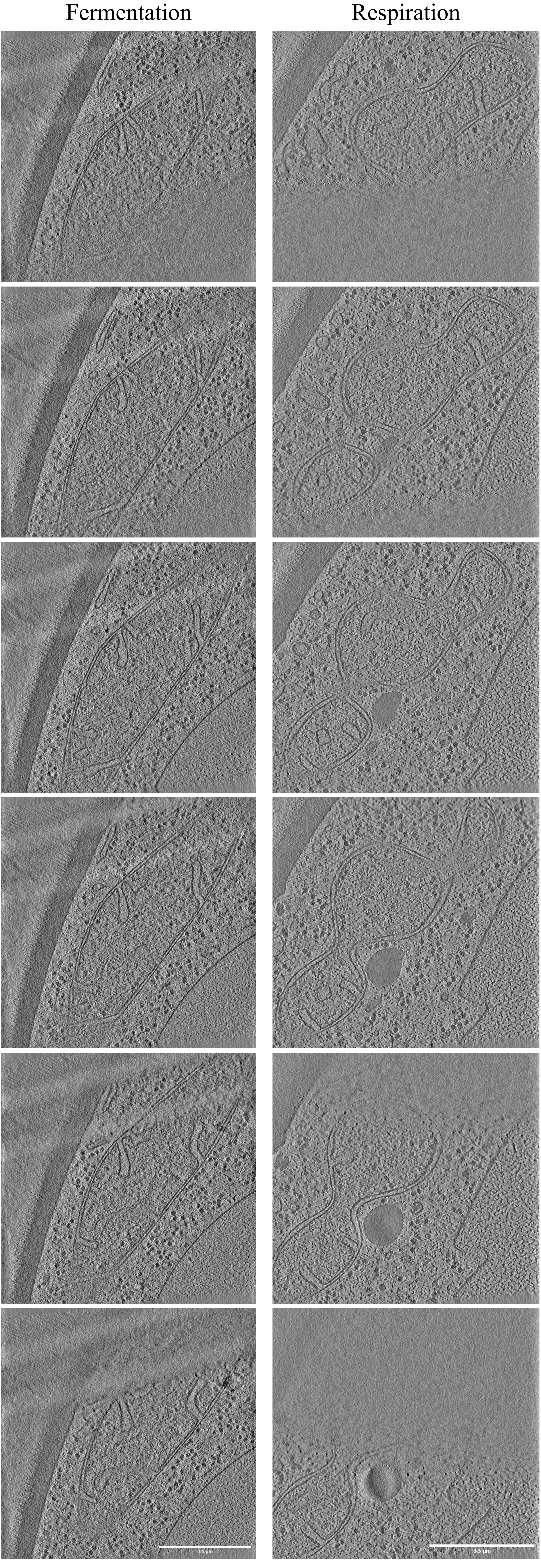

### Extended Data Fig. 6

Singh M. K., et al._Extended Data Figure 6

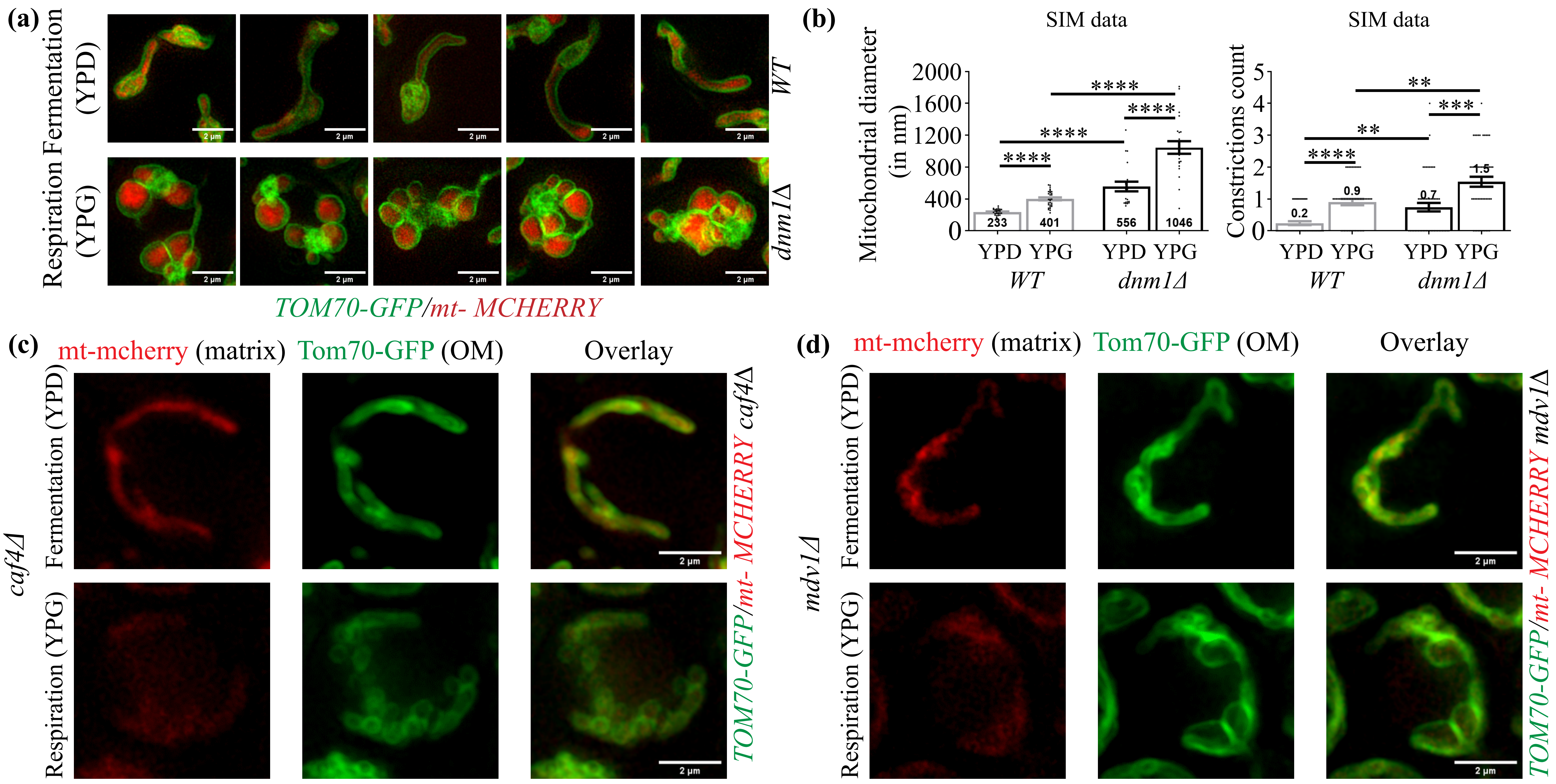

### Extended Data Fig. 7

Singh M. K., et al._Extended Data Figure 7

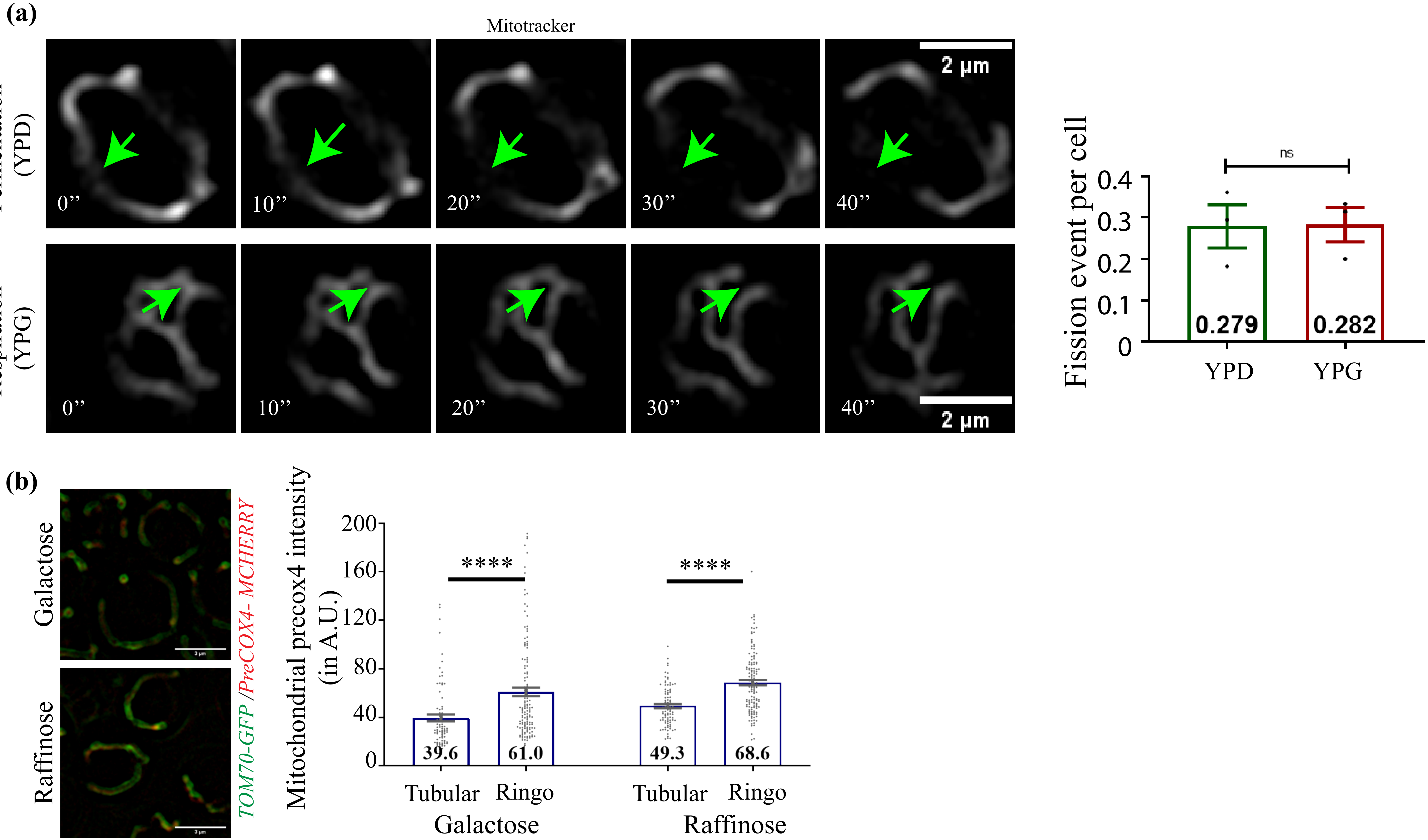

### Extended Data Fig. 8

Singh M. K., et al._Extended Data Figure 8

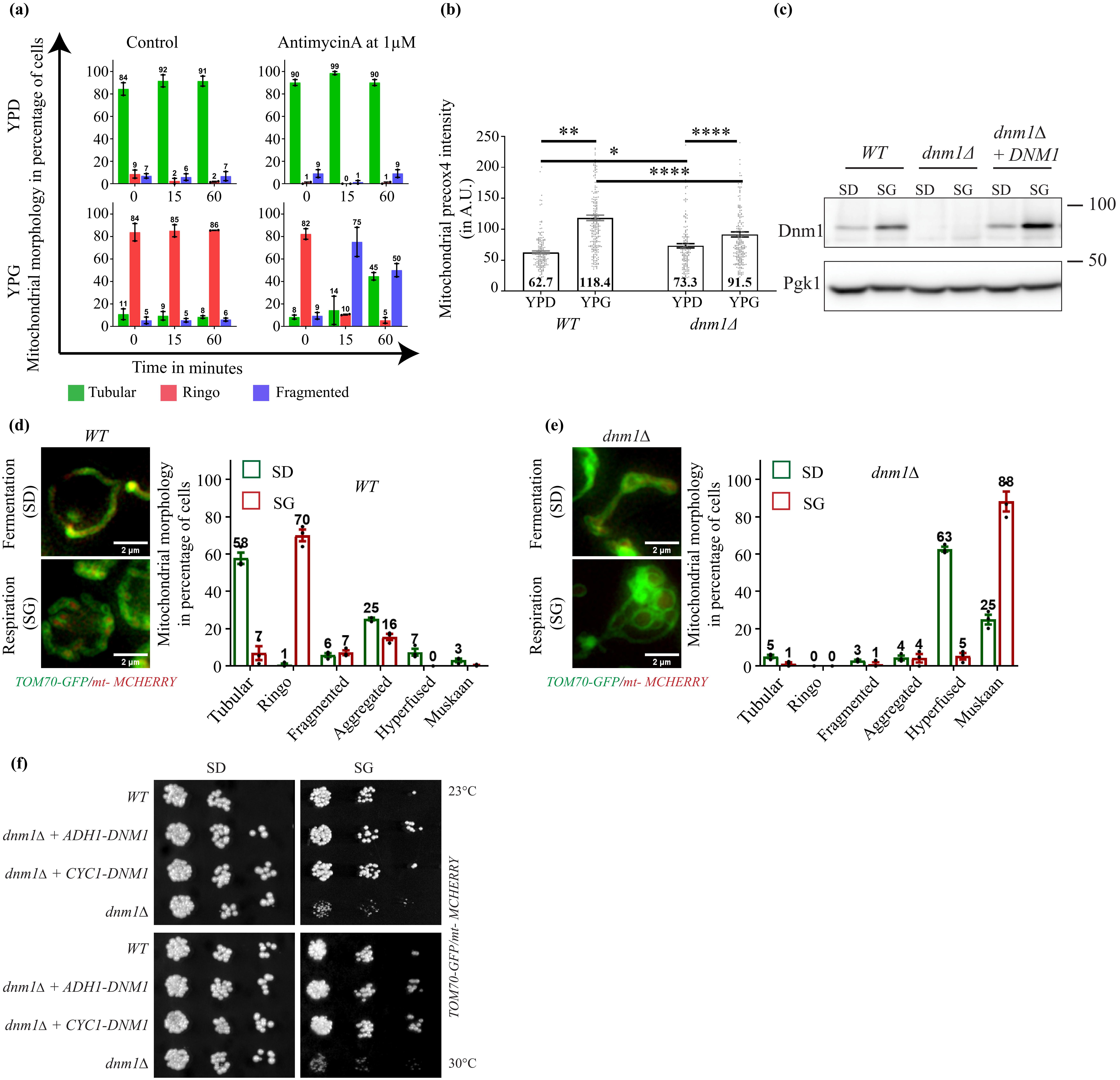
